# BAF perturbation heightens cancer cell dependence on CDK12-driven transcription elongation by RNA polymerase II

**DOI:** 10.64898/2026.07.31.742024

**Authors:** Monika Mačakova, Cassidy Danyko, Stefan Oberlin, Tam Binh V. Bui, Otto Kauko, Houqing Yin, Yuqing Bai, Caroline C. Friedel, Heidi M. Haikala, Annina Vihervaara, Jennifer M. Rosenbluth, Michael T. McManus, Steven Henikoff, Matjaž Barborič

**Author notes:** Equal contribution (Cassidy Danyko, Stefan Oberlin).

## Abstract

CDK12 facilitates transcriptional elongation and processivity by RNA polymerase II (Pol II). Although it has emerged as a promising actor and target in cancer, better understanding of CDK12 gene transcription control could inform the design of novel anti-cancer strategies. Here, we identify a co-dependency between CDK12 and the BAF chromatin-remodeling complex in triple-negative breast cancer (TNBC). Genome-scale CRISPR interference screening revealed multiple BAF subunits as strong dependencies upon CDK12 inhibition. In turn, pharmacological co-targeting of CDK12 and BAF ATPase synergistically reduced the viability of multiple TNBC models. Mechanistically, the co-inhibition attenuated di-phosphorylation of the Pol II C-terminal domain at Serine-2 and Serine-5 and depleted canonical pre-mRNA 3′-end processing factors from chromatin, driving accumulation of chromatin-associated Pol II, upstream of apoptosis. Whereas BAF inhibition rapidly reduced transcription-coupled DNA accessibility, CDK12 inhibition imposed a gene length-biased transcriptional defect exacerbated by BAF deficiency. In turn, co-inhibited cells, marked by synergistic induction of *MYC* and repression of long cell-cycle and mitotic genes, showed a failure of DNA synthesis and mitotic entry, culminating in MYC-mediated apoptosis. Together, our findings reveal a regulatory axis in which impaired Pol II elongation creates a heightened requirement for BAF-dependent chromatin remodeling. This discovered co-dependency provides a rationale for co-targeting the CDK12-BAF axis in transcriptionally addicted cancers.

## INTRODUCTION

Gene transcription by RNA polymerase II (Pol II) shapes cell identity, function, and fate, and is therefore tightly regulated. A central actor in this process is the C-terminal domain (CTD) of the RPB1 subunit of Pol II, a platform whose post-translational marks are written and erased by kinases and phosphatases to steer Pol II activity, precursor mRNA (pre-mRNA) maturation and chromatin modification through the transcription cycle.^1^ Although over one hundred human kinases may directly phosphorylate the CTD Tyr^1^Ser^2^Pro^3^Thr^4^Ser^5^Pro^6^Ser^7^ heptad repeats at signal-responsive genes,^2^ the transcriptional cyclin-dependent kinases (tCDKs) function as the core circuitry controlling distinct steps of Pol II transcription genome-wide.^1,3^ During transcription initiation, CDK7 binds Cyclin H and Mat1 to phosphorylate the CTD on Ser5 and Ser7 residues and promote promoter escape. At the same time, CDK7 enables promoter-proximal pausing of Pol II by stimulating the binding of DRB-sensitivity inducing factor (DSIF) and negative elongation factor (NELF) with Pol II, arresting it in an inactive conformation.^3,4^ The release of Pol II from pausing is triggered by P-TEFb, composed of CDK9 and Cyclin T1 or T2, which as part of the Super elongation complex (SEC) catalyzes di-phosphorylation of the CTD at Ser2 and Ser5 residues (CTD Ser2P-Ser5P) as well as multiple positions within both pause-inducing factors.^5,6^ These events convert DSIF, the SPT4-SPT5 heterodimer, into a positive elongation factor, dissociate NELF from chromatin, and dock the Polymerase associated factor complex (PAF1C) and SPT6 onto Pol II, enabling transcription elongation to resume in a processive manner.^7^ CDK11, which binds Cyclin L1 or L2 and SAP30BP and acts directly in pre-mRNA splicing, has also been reported to stimulate the Pol II pause-release.^8,9^ Finally, CDK12 and CDK13 bind Cyclin K to sustain CTD Ser2P-Ser5P levels, promoting Pol II elongation rate and processivity past intronic polyadenylation sites and through distal exons toward gene 3′ ends.^6,10–12^

Cancer cells depend on misregulated gene expression programs to proliferate and resist cell death. These can be driven not only by specific oncogenic transcription factors (TFs), but also by general Pol II regulators that amplify transcription broadly while preserving transcriptional fidelity. Accordingly, hypertranscription has been recognized as a hallmark of aggressive human cancers.^13,14^ This transcriptional dependency is exemplified by oncogenic circuits that co-opt Pol II elongation machinery.^5,15^ For example, misregulated transcription driven by the MYC-MAX heterodimer characterizes a broad range of human cancers, including breast, colon, cervix, lung and pancreatic cancer, as well as several types of leukemias and lymphomas.^15–17^ Similarly, mixed lineage leukemia gene (*MLL*)-translocation partners, several of which encode SEC subunits, redirect SEC to aberrant genomic loci to enforce oncogenic gene regulation in pediatric leukemias.^5^ Accordingly, components of the Pol II transcription cycle, including transcriptional CDKs and elongation factors, have emerged as druggable vulnerabilities in cancer.^3,15^

Among tCDKs, CDK12 and its paralogue CDK13 are particularly relevant in cancer. The role of CDK12 in stimulating productive Pol II elongation rate and processivity becomes critical at long and complex genes, including those involved in the homologous recombination (HR) DNA repair pathway, DNA replication and cell-cycle.^10,11,18–20^ Consistent with this role, genetic or pharmacological disruption of CDK12 induces genomic instability. Inactivating *CDK12* alterations occur in several cancers, including high-grade serous ovarian carcinoma and metastatic castration-resistant prostate cancer, where CDK12-deficient tumors are characterized by HR deficiency reminiscent of BRCAness phenotype, large tandem duplications, and increased neoantigen burden.^18,21–25^ This connection between CDK12 loss and impaired DNA repair provided a rationale for exploiting CDK12-deficient tumors with DNA damage response therapies, including PARP inhibitors.^26–28^ By contrast, disruption of CDK13 impairs PAXT-mediated nuclear RNA surveillance, promoting oncogenic gene-expression.^29^ In other contexts, CDK12 and CDK13 can promote oncogenesis, highlighting their context-dependent role in cancer.^30–33^

Selective co-targeting of CDK12 and CDK13 (hereafter referred to as CDK12 for simplicity) by the dual, covalent inhibitor THZ531 has been pivotal in dissecting the roles of these tCDKs in gene regulation.^34^ Cyclin K degraders such as SR-4835 later expanded the therapeutic exploration of CDK12 targeting in cancer models, both as single agents and in combination with other targeted therapies or immunotherapies.^28,35–39^ However, the regulatory dependencies that determine sensitivity to CDK12 inhibition remain incompletely understood, leaving gene-control mechanisms and therapeutic opportunities undiscovered.

To systematically identify targetable vulnerabilities associated with CDK12 inhibition, we performed a genome-scale CRISPR interference (CRISPRi) screen in THZ531-treated triple-negative breast cancer cells. We identified a functional dependency of CDK12-inhibited cells on the BRG1/BRM-associated factor (BAF) chromatin-remodeling complex, also known as the mammalian SWI/SNF. BAF complexes are large, multi-subunit ATP-dependent chromatin remodelers that use ATP hydrolysis to reposition or evict nucleosomes, shaping DNA accessibility and TF occupancy across promoters and enhancers.^40–42^ While genes that encode subunits of BAF chromatin remodeling complexes are mutated in over 20% of cancers^42,43^, oncogenic TFs can hijack wild-type (WT) BAF complexes to promote oncogenesis^44–46^, spurring a major interest in developing BAF-targeted therapies.^42^ Here, we define a functional relationship between CDK12-dependent transcription elongation and BAF-dependent chromatin remodeling. Genetic or pharmacological BAF perturbation sensitizes TNBC cells to CDK12 inhibition, producing synergistic loss of viability across multiple models. Mechanistically, combined CDK12 and BAF inhibition disrupts Pol II elongation and transcription-coupled DNA accessibility at regulatory loci. The resulting gene length-biased transcriptional perturbation, characterized by synergistic induction of *MYC* and repression of long cell-cycle and mitotic genes, drives a failure in DNA synthesis and mitotic entry, culminating in MYC-mediated apoptosis. Together, our findings identify BAF-dependent chromatin remodeling as a targetable vulnerability in CDK12-inhibited cancer cells,

## RESULTS

### CRISPRi screen identifies a functional dependency of CDK12-inhibited cells on the BAF chromatin-remodeling complex

To identify genes required for cancer cell survival upon CDK12/13 inhibition, we performed a genome-scale CRISPR interference (CRISPRi) screen in the TNBC cell line MDA-MB-231 (Fig. 1A). To do so, we generated a monoclonal MDA-MB-231 CRISPRi line stably expressing ZIM3 KRAB-dCas9,^47^ and selected a clone with efficient sgRNA-mediated silencing of cell-surface antigen CD146 by flow cytometry (Fig. S1A). We transduced the cells at low multiplicity of infection (MOI = 0.3) with the Dolcetto sgRNA library which targets each protein-coding gene with six independent sgRNAs plus 500 non-targeting control sgRNAs.^48^ Following selection, we cultured the cells for 18 days in the presence of either DMSO or THZ531 at its IC50 concentration of 110 nM (Fig. 1A, Fig. S1B).

**Figure 1.**
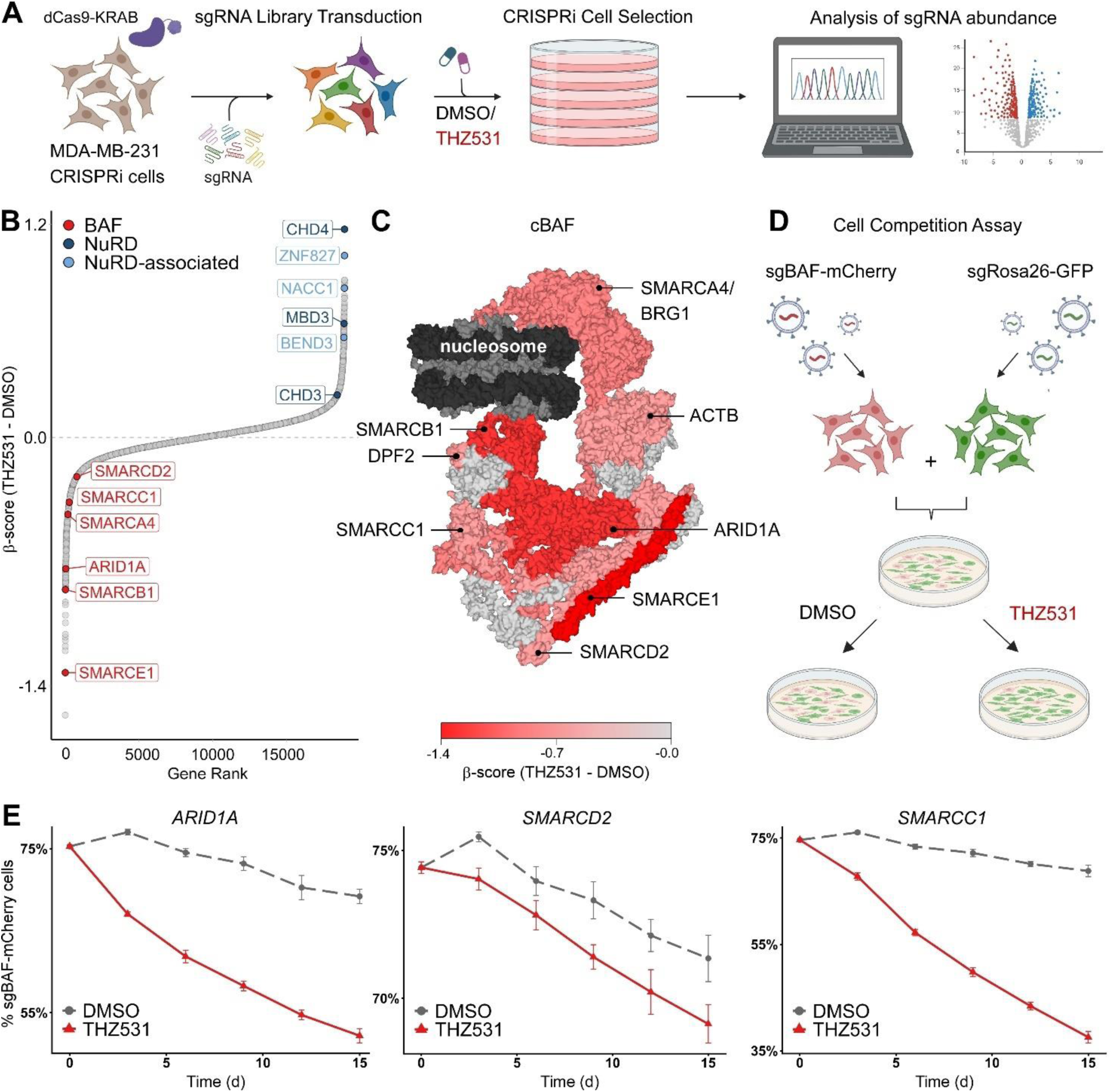
Genetic perturbation of the BAF chromatin-remodeling complex creates a vulnerability in CDK12-inhibited MDA-MB-231 cells. **A)** Schematic of the genome-wide CRISPRi screen in MDA-MB-231 CRISPRi cells. Cells were transduced with the Dolcetto sgRNA library, treated with DMSO or THZ531 (110 nM) for 18 days, and surviving cells were analyzed by sgRNA sequencing. **B)** Waterfall plot representing gene-level results of the CRISPRi screen. BAF, NuRD, and NuRD-associated genes of which sgRNAs were enriched or depleted in THZ531-treated cells relative to DMSO are highlighted. **C)** Structural schematic of the BAF complex with screen-identified subunits highlighted. Red intensity indicates THZ531-selective depletion of sgRNAs targeting the indicated BAF subunits. Remodeled histone octamer and DNA are shown in dark gray and black, respectively. Structure: PDBDEV_00000056.^121^ **D)** Schematic of the fluorescence-based cell competition assay. **E)** Relative abundance of mCherry-positive CRISPRi cells targeting the indicated BAF subunits during competition with GFP-positive sgRosa26 control cells under DMSO or THZ531 (110 nM) treatment for two weeks. Data are shown as mean ± s.e.m. (n = 2).

Quality-control metrics confirmed high sgRNA representation, even library distribution across conditions, and depletion of essential genes (Fig. S1C,D). MAGeCK analysis^49,50^ identified multiple subunits of the canonical BAF complex as dependencies upon CDK12 inhibition (Fig. 1B,C, Fig. S1E). These included the ATPase SMARCA4/BRG1 and the core or regulatory subunits ARID1A, SMARCB1, SMARCC1, SMARCE1 and SMARCD2. In contrast, silencing subunits of the repressive Nucleosome Remodeling and Deacetylase (NuRD) complex, including CHD4, CHD3 and MBD3, conferred resistance to CDK12 inhibition (Fig. 1B). That BAF and NuRD perturbation elicited opposing phenotypes is consistent with their antagonistic roles in gene regulation.^51–53^

To validate these findings, we performed cell competition assays using MDA-MB-231 CRISPRi cells. We mixed mCherry-positive cells expressing sgRNAs targeting the BAF *ARID1A*, *SMARCD2*, or *SMARCC1* with GFP-positive cells expressing a *Rosa26* safe-targeting sgRNA, treated the mixed cultures with DMSO or THZ531, and monitored population dynamics by flow cytometry over 14 days (Fig. 1D). BAF-targeted cells remained largely stable under DMSO but were progressively outcompeted by *Rosa26* control cells under THZ531, confirming that BAF perturbation sensitizes cells to CDK12 inhibition (Fig. 1E). Conversely, targeting the NuRD *CHD4* and *MBD3*, or the NuRD-associated *ZNF827*, *NACC1*, and *BEND3*, showed the opposite behavior (Fig. S1F). These competition assays independently validated the screen findings for BAF and NuRD. Together, these data show that BAF perturbation creates a vulnerability in CDK12-inhibited TNBC cells.

### Pharmacological co-targeting of CDK12 and BAF decreases viability of TNBC models in a synergistic manner

Given the identified functional relationship, we hypothesized that pharmacological co-targeting of CDK12 and BAF using highly selective inhibitors would decrease TNBC cell viability. To test this, we exposed MDA-MB-231 cells to combinatorial titrations of THZ531 and the BRG1/BRM ATPase inhibitor FHT-1015^54^, and quantified cytotoxicity and viability after three days. Compared with either agent alone, the co-targeting of CDK12 and BAF led to increased cytotoxicity and reduced viability (Fig. 2A). As revealed by the Bliss synergy scores calculated by Synergyfinder,^55^ these effects were highly synergistic across a broad range of inhibitor concentrations, reaching a maximum Bliss score of 68 (Fig. 2A, bottom), well above the commonly used synergy threshold of ten. We recapitulated these synergistic effects by replacing one arm of the CDK12-BAF combination with a distinct compound: on the BAF arm, the dual BRG1/BRM degrader ACBI1^56^ (Fig. 2B), and on the CDK12 arm, the Cyclin K degrader SR-4835^36^ (Fig. S2A).

**Figure 2.**
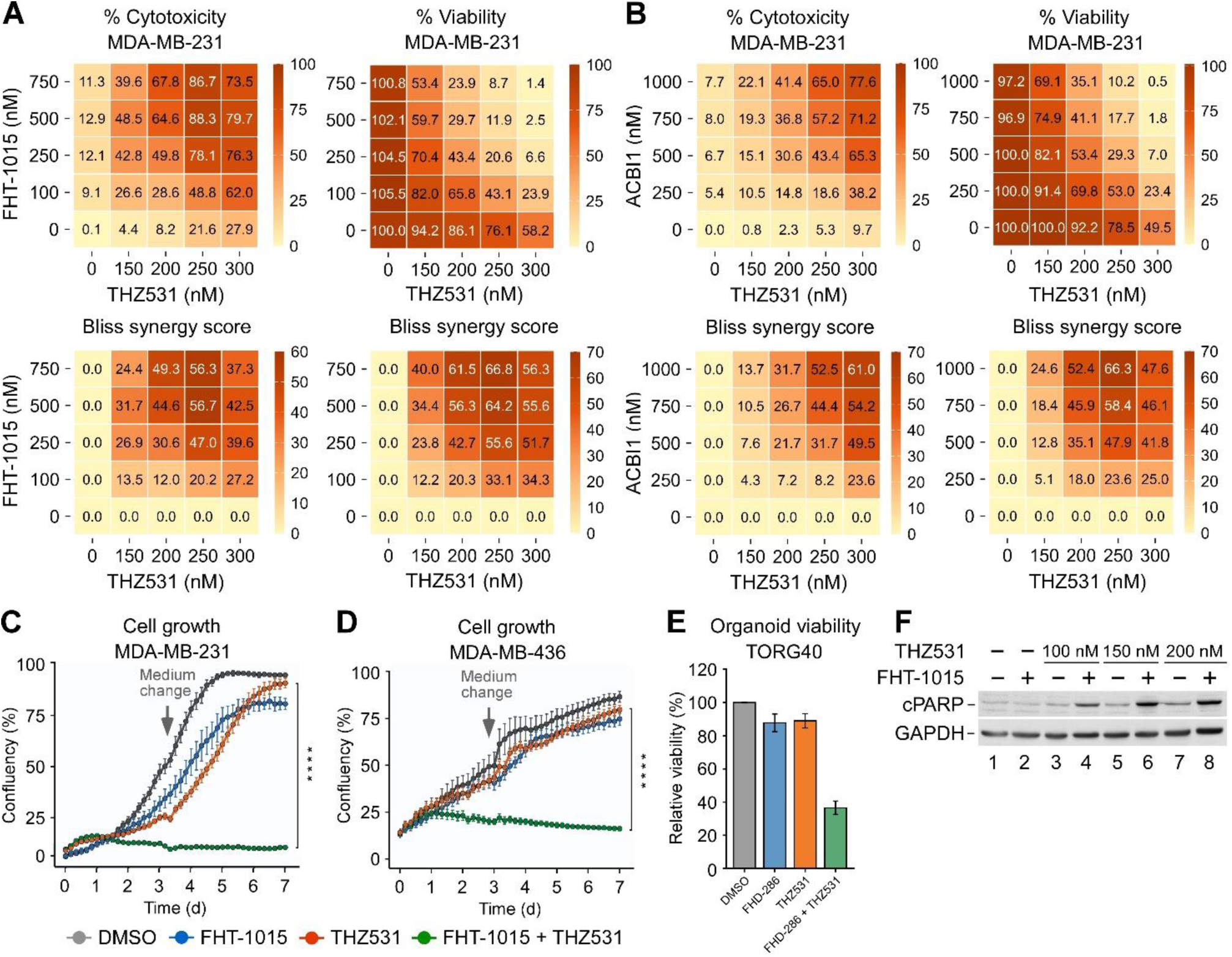
Pharmacological co-targeting of CDK12 and BAF synergizes in decreasing viability of TNBC models. **A,B)** Cytotoxicity and viability matrices with combinatorial titrations of FHT-1015 (A) or ACBI1 (B) and THZ531 at indicated concentrations in MDA-MB-231 cells, depicting cytotoxicity and viability (top), and Bliss synergy scores (bottom) at 72 h. Cytotoxicity and viability data are shown as percentages relative to DMSO-treated cells and represent the average of independent experiments (n = 3). **C,D)** Cell growth of MDA-MB-231 (C) and MDA-MB-436 (D) cells treated for seven days with DMSO, FHT-1015 (500 nM), THZ531 (60 nM), and the combination as indicated. Results are presented as % confluency and plotted as the mean ± s.e.m. (n = 3). Arrows indicate the replenishment of medium and drugs at day 3. \*\*\*\**P* < 0.0001, determined by two-way ANOVA using THZ531 and FHT-1015 + THZ531 data sets. **E)** Relative organoid viability treated for 72 h with DMSO, FHD-286 (1000 nM), THZ531 (50 nM), and the combination as indicated. Data are shown relative to DMSO-treated organoid and plotted as the mean ± s.e.m. (n = 3). **F)** Immunoblot analysis of cPARP and GAPDH in whole cell extracts of MDA-MB-231 cells treated for 24 h with FHT-1015 (750 nM), increasing concentrations of THZ531, and the combinations as indicated.

To provide complementary evidence for the synergistic loss of cell viability by the CDK12 and BAF co-targeting, we exposed MDA-MB-231 cells to the same drug combinations and monitored their growth over seven days using live-cell imaging. In contrast to the single-agent treatments, combined perturbation of CDK12 and BAF attenuated cell growth in a synergistic manner (Fig. 2C and Fig. S2B,D). We conducted the co-inhibition studies using additional TNBC cell lines, including MDA-MB-436 and HCC1395 cells, which also showed pronounced and synergistic sensitivity to CDK12 and BAF co-inhibition (Fig. 2D and Fig. S2C,E,F).

Beyond these monolayer cell line experiments, we tested the potency of combined CDK12 and BAF inhibition in a previously established and characterized organoid model derived from a TNBC patient.^57^ We selected TORG40 as a model that lacks mutations in BAF subunit genes. Here, we combined THZ531 with FHD-286, a dual BRG1/BRM ATPase inhibitor employed previously in pre-clinical and clinical studies.^58–60^ Whereas either inhibitor alone affected the organoids only marginally, their combination reduced viability markedly (Fig. 2E). These findings extend the suppressive activity of CDK12–BAF co-targeting to a patient-derived TNBC model.

We nest asked whether the combined perturbation induced cell death by apoptosis. We conducted immunoblot analyses to monitor levels of the apoptotic marker caspase 3-cleaved poly(ADP-ribose) polymerase (cPARP) in whole cell extracts of MDA-MB-231 cells exposed to inhibitor concentrations that, when combined, elicited synergistic loss of viability in the assays above. Single-agent BAF inhibition with FHT-1015 or CDK12 inhibition with increasing THZ531 elevated PARP cleavage only modestly, whereas co-targeting increased cPARP synergistically (Fig. 2F). Together, these results establish that pharmacological co-targeting of CDK12 and BAF suppresses TNBC cell growth and promotes apoptosis.

### Combined CDK12 and BAF inhibition attenuates Pol II CTD Ser2P-Ser5P di-phosphorylation and depletes pre-mRNA 3′-end processing factors from chromatin

The kinase activity of CDK12 stimulates transcription elongation and processivity by Pol II. Therein, the CDC73 subunit of PAF1C allosterically activates CDK12 to catalyze di-phosphorylation of the Pol II CTD at Serine-2 and Serine-5 residues (CTD Ser2P-Ser5P).^6^ To investigate the molecular basis for the synergistic loss of cancer cell viability upon CDK12-BAF co-targeting, we determined the levels of the CTD Ser2P-Ser5P elongation mark and total Pol II in chromatin fractions isolated from MDA-MB-231 cells exposed to synergistic inhibitor concentrations in isolation or combination across an eight-hour time course. We also monitored the levels of selected subunits of CDK12 and BAF, CDC73 of PAF1C, and histone H3 as a chromatin marker. Unlike the BAF inhibitor FHT-1015, a sub-lethal dose of CDK12 inhibitor THZ531 alone resulted in a modest decrease of CTD Ser2P-Ser5P. Critically, the combination attenuated this elongation mark more than either agent alone by 8 h, while the unphosphorylated form of Pol II accumulated, suggesting deficient transcription elongation (Fig. 3A and Fig. S3A). In contrast, levels of the other proteins tested remained stable. We observed a similar defect in co-targeted cells when we substituted the BAF ATPase inhibitor with the degrader ACBI1 (Fig. 3B). There, levels of BRG1 and SMARCB1 subunits showed a progressive decrease as expected.

**Figure 3.**
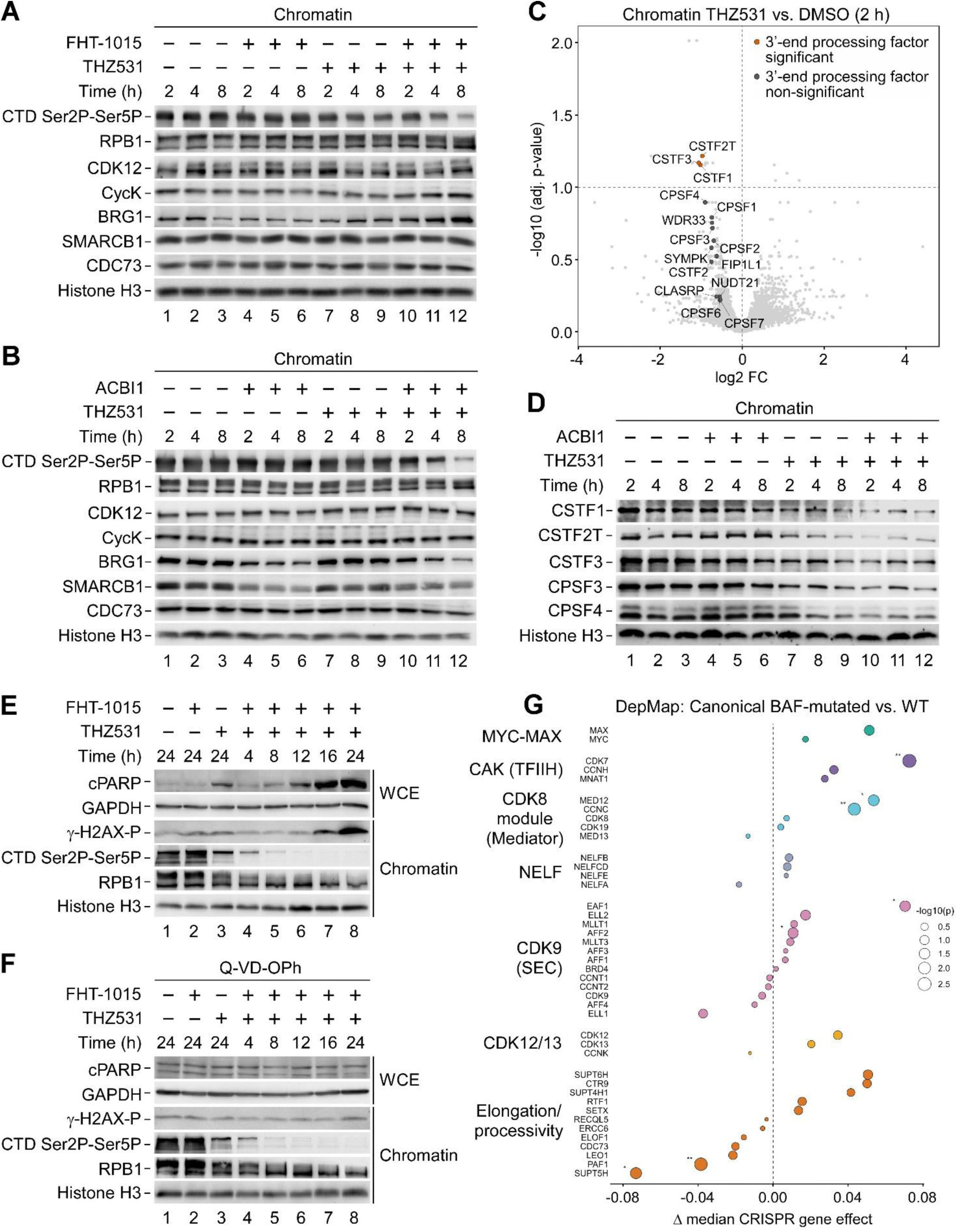
Combined CDK12 and BAF inhibition impairs CTD Ser2P-Ser5P di-phosphorylation in synergistic and apoptosis-independent manner. **A,B)** Immunoblot analysis of the indicated proteins in chromatin fractions of MDA-MB-231 cells treated with DMSO (–), FHT-1015 (750 nM; A), ACBI1 (1000 nM; B), THZ531 (150 nM), and the combinations for 2, 4, and 8 h as indicated. **C)** Volcano plot showing log2 FC values (adjusted *P* ≤ 0.1) of 3’-end processing factor levels in chromatin fraction of MDA-MB-231 cells treated with THZ531 (400 nM) for 2 h, relative to DMSO (n = 4). Significant and non-significant factors are shown in red and dark gray, respectively; other proteins are shown in light gray. **D)** Immunoblot analysis of the indicated proteins in chromatin fractions of MDA-MB-231 cells treated with DMSO (–), ACBI1 (1000 nM), THZ531 (150 nM), and the combinations for 2, 4, and 8 h as indicated. **E,F)** Time-course immunoblot analysis of the indicated proteins in whole-cell extracts (WCE) and chromatin fractions of MDA-MB-231 cells treated with DMSO (–), FHT-1015 (750 nM), THZ531 (150 nM), and the combinations for the indicated durations with or without Q-VD-OPh (25 μM) as indicated. **G)** DepMap CRISPR dependency analysis comparing canonical BAF-mutated cancer models (n = 243) with BAF-WT/non-mutated models (n = 949). Transcriptional regulators were grouped according to their roles across the Pol II transcription cycle. Ä median gene-effect scores indicate differential dependency, with negative values reflecting stronger dependency in BAF-mutated models. Point size indicates −log10(nominal P); asterisks indicate nominal Wilcoxon rank-sum significance (*P < 0.05, **P < 0.01).

To provide an explanation for increased accumulation of unphosphorylated Pol II on chromatin upon combined CDK12 and BAF inhibition, we performed an unbiased proteomic analysis of chromatin fractions isolated from MDA-MB-231 cells. Strikingly, CDK12 inhibition elicited coordinated reduction in chromatin association of multiple components of the canonical pre-mRNA 3′-end processing machinery (Fig. 3C), which produces polyadenylated transcripts and couples 3’-end pre-mRNA cleavage to Pol II termination.^61,62^ The cleavage stimulation factor (CSTF) complex, comprising CSTF1/CstF50, CSTF2/CstF64, CSTF2T/τCstF64 and CSTF3/CstF77, recognizes U/GU-rich sequences downstream of polyadenylation signal (PAS) sequence and cooperates with the cleavage and polyadenylation specificity factor (CPSF) complex to promote pre-mRNA cleavage.^61^ CSTF1, CSTF2T and CSTF3 were significantly depleted from chromatin following CDK12 inhibition, while multiple CPSF components, including CPSF1/CPSF160, CPSF4/CPSF30, WDR33, FIP1L1/hFip1, and mammalian cleavage factor subunits CPSF2/CPSF100, CPSF3/CPSF73 and Symplekin, showed a similar trend that did not reach our significance threshold (Fig. 3C). We validated selected components of both complexes by independent chromatin fractionation and immunoblotting, confirming depletion of CSTF1/CstF50, CSTF2T/τCstF64, CSTF3/CstF77, CPSF3/CPSF73 and CPSF4/CPSF30 from chromatin following CDK12 inhibition in an eight-hour time course experiment. Notably, while BAF perturbation exerted no effect, it further reduced the chromatin depletion of these factors elicited by CDK12 inhibition (Fig. 3D and Fig. S3B). These findings suggest that the chromatin depletion of CSTF and CPSF contributes to altered pre-mRNA 3′-end processing and Pol II termination, contributing to accumulation of Pol II on chromatin.

We next asked whether the Pol II CTD di-phosphorylation defect preceded apoptotic execution in CDK12–BAF co-targeted MDA-MB-231 cells. In a 24 h time-course experiment, loss of chromatin CTD Ser2P-Ser5P preceded PARP cleavage (Fig. 3E). Because CDK12 inhibition elicits DNA damage, we also monitored γ-H2AX Ser139 phosphorylation (γ-H2AX-P), a marker of DNA double-strand breaks. Chromatin levels of γ-H2AX-P accumulated strongly at 24 h post-combination treatment, but with delayed kinetics relative to PARP cleavage. Pan-caspase inhibitor Q-VD-OPh blocked PARP cleavage and γ-H2AX-P accumulation (Fig. 3F), indicating that caspase activity is required for the γ-H2AX-P accumulation, placing DNA-damage signaling downstream of apoptosis in this setting.

Since combined CDK12 and BAF inhibition resulted in a Pol II elongation defect, we hypothesized that BAF-perturbed cancers might be dependent broadly on Pol II elongation machinery. To address this, we analyzed DepMap CRISPR dependency data by comparing the reliance of canonical BAF-mutated cancer models and BAF-WT/non-mutated models on regulators controlling distinct phases of the Pol II transcription cycle (n = 243 versus 949). Consistent with this prediction, canonical BAF-mutated models showed preferential dependency on specific Pol II elongation and processivity factors, most notably SPT5 of DSIF, PAF1, LEO1 and CDC73 of PAF1C, and ELL1 of the CDK9-containing SEC (Fig. 3G and Fig. S3C). DepMap subgroup analysis revealed the strongest SPT5 dependency in ARID1A/B-mutated cell lines, consistent with the role of SPT5 in maintaining the enhancer landscape by direct interaction with BAF subunits BRG1, SMARCC2, and ARID1A^63^ (Fig. S3D). Furthermore, SMARCC1/2-mutated cell lines showed particularly strong dependency on PAF1 (Fig. S3E). Finally, although our pan-canonical BAF-mutant analysis did not reveal a strong dependency on the CDK12/13–Cyclin K module itself, SMARCC1/2-mutated models showed increased dependency on Cyclin K (Fig. S3E), consistent with the loss of cell viability we observed by pharmacological co-targeting of CDK12 or Cyclin K and BAF.

Together, we conclude that combined CDK12 and BAF inhibition causes an early, apoptosis-independent loss of CTD di-phosphorylation, indicative of perturbed transcription elongation. Given that 3’-end pre-mRNA cleavage acts as a trigger for Pol II termination,^62^ the concurrent accumulation of chromatin-associated Pol II and depletion of canonical 3’-end processing factors in co-inhibited cells suggests impaired coupling of elongation-deficient Pol II to productive 3′-end processing and termination. Finally, our DepMap analysis findings raise a possibility that dependence on Pol II elongation machinery may represent a broader transcriptional vulnerability in BAF-mutated cancers.

### CDK12 and BAF inhibition decrease BAF occupancy and transcription-coupled DNA accessibility

Having established that the CDK12-BAF co-targeting synergistically decreases cell viability and Pol II elongation, we next used genomic approaches to probe the functional relationship of CDK12 and BAF. Although the consequences of BAF perturbation in cancer have been studied extensively,^42,43^ less is known about how BAF occupancy and activity intersect with the Pol II transcription machinery. We hypothesized that CDK12 and BAF co-occupy gene promoters and enhancers, and that inhibiting them alters their chromatin occupancy and transcription-coupled DNA accessibility.

We first used Cleavage Under Targets and Tagmentation (CUT&Tag) to map genome-wide CDK12 and BAF occupancy *in situ*.^64^ We applied CUT&Tag by using antibodies targeting CDK12 and BRG1, the catalytic subunit of BAF responsible for ATP-dependent nucleosome remodeling. Unlike BRG1, CDK12 does not bind DNA directly. CDK12 CUT&Tag therefore likely captures only the most enriched, stable CDK12-occupied sites. The top 20,000 peaks for each factor were annotated to cis-regulatory elements, revealing substantial BRG1 and CDK12 occupancy at both promoters and enhancers (Fig. 4A and Fig. S4A,B). We then classified promoter and enhancer elements as CDK12-BRG1 co-occupied, BRG1-only, or CDK12-only. CDK12 and BRG1 co-occupied 44% of promoter-associated and 19% of enhancer-associated elements, while approximately half of enhancer-associated elements were occupied by BRG1 alone (Fig. S4C).

**Figure 4.**
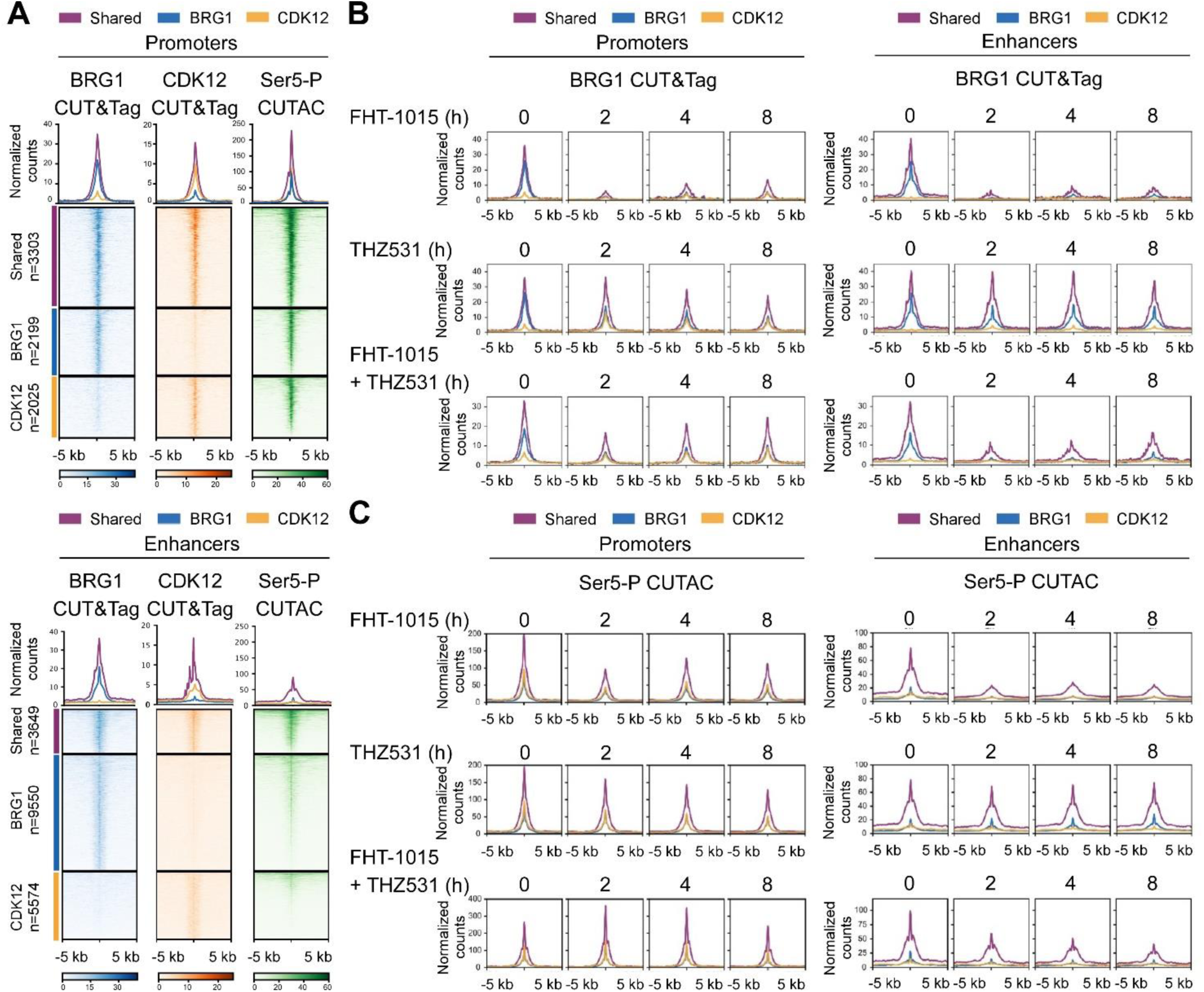
Catalytic activities of CDK12 and BAF promote their chromatin occupancy and transcription-coupled DNA accessibility. **A)** Average profiles (above) and heatmaps (below) of BRG1 CUT&Tag, CDK12 CUT&Tag, and Ser5-P CUTAC signal at BRG1 and CDK12 co-occupied (shared), BRG1-only, and CDK12-only promoters and enhancers in MDA-MB-231 cells (n = 2). Average profiles and heatmaps show normalized signal across the indicated regulatory-element classes. Promoter and enhancer profiles are centered on transcription start sites and enhancer mid-points, respectively. **B,C)** Average profiles of BRG1 CUT&Tag (B) and Ser5-P CUTAC (C) signal at BRG1 and CDK12 co-occupied (shared), BRG1-only, and CDK12-only promoters and enhancers in MDA-MB-231 cells treated with FHT-1015 (1 μM), THZ531 (400 nM), and the combination for 2, 4, and 8 h as indicated (n = 2). Average profiles show normalized signal across the indicated regulatory-element classes. Promoter and enhancer profiles are centered on transcription start sites and enhancer mid-points, respectively.

We next examined whether perturbing CDK12 and BAF using THZ531 and FHT-1015, respectively, affects their chromatin occupancy over an eight-hour period. As expected, inhibiting the BAF ATPase strongly reduced BRG1 occupancy at both promoters and enhancers (Fig. 4B and Fig. S4D). Consistently, chromatin proteomic analysis revealed profound BAF perturbation upon FHT-1015 treatment: in addition to BRG1, the ATPase inhibition depleted multiple BAF subunits from chromatin, which relocated to the nucleoplasm (Fig. S4E). BAF inhibition also reduced CDK12 occupancy (Fig. S4F), whereas CDK12 inhibition gradually reduced BRG1 occupancy, particularly at co-occupied promoters and enhancers (Fig. 4B and Fig. S4D), consistent with an interdependent relationship between active CDK12 and the BAF complex on chromatin.

To extend these findings, we asked whether CDK12 and BAF activities regulate DNA accessibility at regulatory sites marked by transcriptionally engaged Pol II. We therefore performed low-salt tagmentation CUT&Tag in MDA-MB-231 cells using an antibody against Ser5-phosphorylated Pol II. This approach, referred to as Ser5-P Cleavage Under Targeted Accessible Chromatin (Ser5-P CUTAC), identifies accessible chromatin associated with Ser5-phosphorylated Pol II and provides high-resolution maps of transcription-coupled regulatory sites, including active promoters and enhancers.^65,66^

When aligned with regulatory elements occupied by CDK12 and BRG1, the co-occupied sites showed markedly higher Ser5-P CUTAC signal than those occupied by either factor alone (Fig. 4A), consistent with the cooperative function of CDK12 and BAF at highly accessible, Pol II-engaged chromatin. CDK12 inhibition resulted in a gradual decrease of transcription-associated chromatin at promoters but not at enhancers, broadly paralleling the reduced BRG1 occupancy at co-occupied regulatory sites (Fig. 4C and Fig. S4G). In contrast to enhancers, the prompt loss of BRG1 occupancy in BAF-inhibited cells did not translate into sustained loss of Ser5-P CUTAC signal at promoters (Fig. 4C and Fig. S4G).

Finally, combined CDK12 and BAF inhibition reduced both BRG1 occupancy and transcription-coupled DNA accessibility at enhancers, resembling the effect of BAF inhibition alone (Fig. 4B,C, Fig. S4D,G). At promoters, however, the co-inhibition produced a Ser5-P CUTAC pattern that did not mirror either single perturbation, suggesting that Pol II pausing dynamics upon CDK12 inhibition intersects with compensatory chromatin remodeling mechanisms.^67^

Together, these data indicate that the catalytic activities of CDK12 and BAF support their co-occupancy on chromatin, which marks highly accessible chromatin with engaged Pol II. Furthermore, within the eight-hour window analyzed here, BAF perturbation, alone or combined with CDK12 inhibition, preferentially compromised transcription-coupled DNA accessibility at distal enhancers, whereas promoters remained comparatively resilient, consistent with the idea that promoter accessibility established by BAF can be maintained by promoter-dedicated compensatory mechanisms when BAF activity is impaired.

### BAF inhibition exacerbates gene length-biased transcriptional response in CDK12-inhibited TNBC cells

As changes in CDK12 and BAF occupancy and transcription-coupled DNA accessibility do not necessarily predict final RNA output, we performed mRNA sequencing (mRNA-seq) to ask how CDK12 and BAF perturbation reshaped steady-state gene expression. We isolated poly-A enriched RNA pools from MDA-MB-231 cells treated for eight hours with DMSO, FHT-1015 and THZ531 alone or in combination, a time point before CDK12-BAF co-inhibition triggers apoptosis (Fig. 3C). Based on the reproducible mRNA-seq data (Fig. S5A), we determined differentially expressed genes (adjusted P ≤ 0.001; absolute log2FC ≥ 1). BAF inhibition alone produced a modest transcriptional response. Of the 13,983 mRNA-producing genes in MDA-MB-231 cells, 580 were down-regulated and 249 were up-regulated after eight hours of BAF inhibition (Fig. 5A, left). In contrast, CDK12 inhibition produced a much larger response, altering 34% of genes, about 60% of which were down-regulated (Fig. 5A, middle). Finally, co-inhibition produced the largest perturbation, altering 37% of genes (Fig. 5A, right).

**Figure 5.**
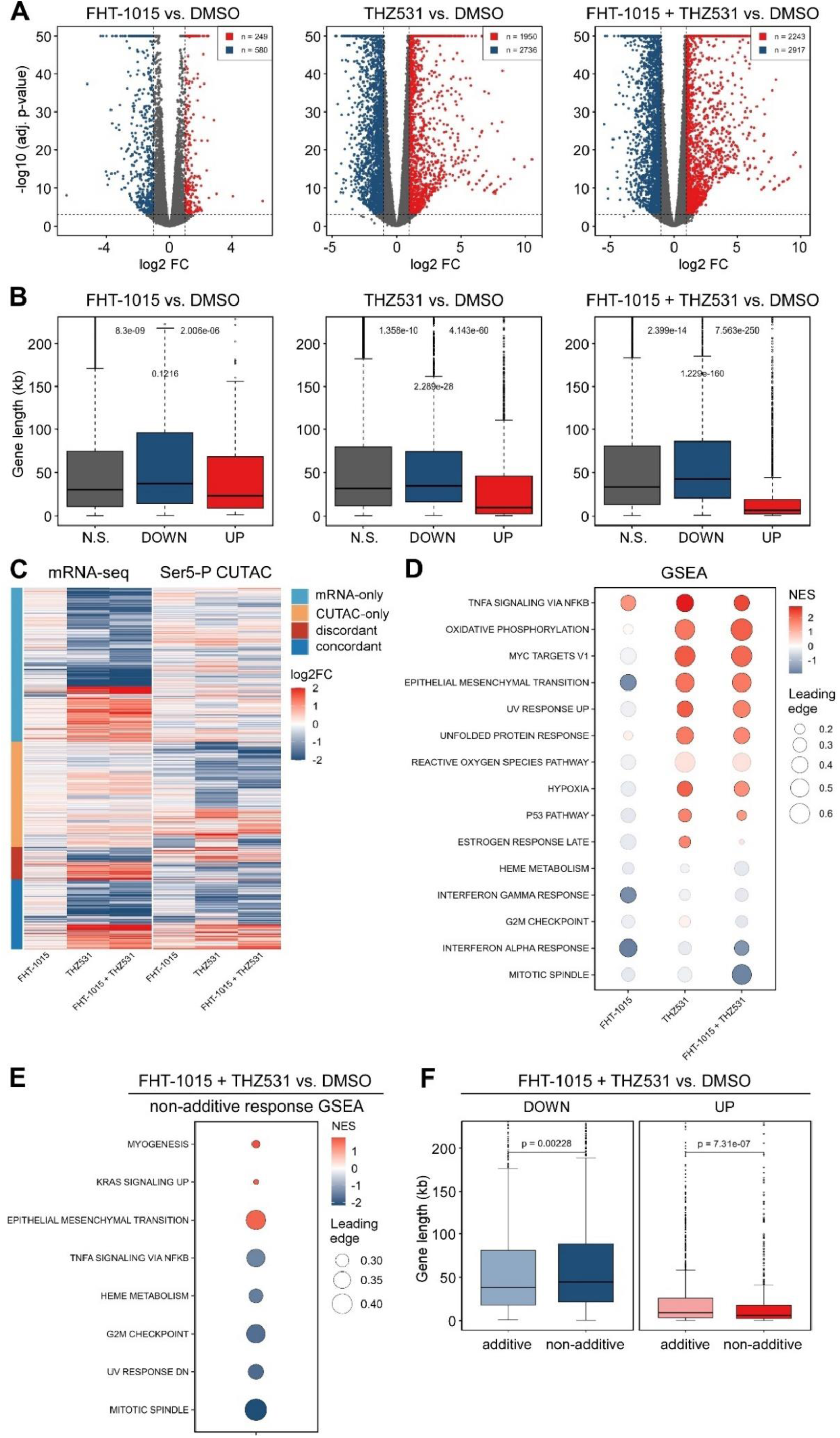
CDK12 and BAF perturbation reshapes gene expression programs in TNBC cells. **A)** Volcano plot representation of differentially expressed genes (*n* = 13 983; *P*-adj ≤ 0.001; Log2FC ≥ 1) from mRNA-seq experiments of MDA-MB-231 cells treated for 8 h with DMSO, FHT-1015 (750 nM), THZ531 (300 nM), and the combination as indicated (n = 3). Red, blue, and gray indicate up-regulated, downregulated, and non-significantly regulated genes, respectively; numbers indicate the significantly regulated genes. **B)** Gene-length distribution of non-significantly regulated (N.S.), down-regulated (DOWN), and up-regulated (UP) protein-coding genes (*n* = 13 983; *P*-adj ≤ 0.001; Log2FC ≥ 1) from mRNA-seq experiments of MDA-MB-231 cells treated for 8 h with DMSO, FHT-1015 (750 nM), THZ531 (300 nM), and the combination as indicated (n = 3). Boxplots show median and interquartile range; P values indicate pairwise Wilcoxon rank-sum tests comparing down-regulated genes against non-significantly or up-regulated genes (top) or between non-significantly and up-regulated genes (middle). **C)** Heatmap comparing mRNA-seq and Ser5-P CUTAC changes of MDA-MB-231 cells treated for 8 h with DMSO, FHT-1015 (750 nM), THZ531 (300 nM), and the combination as indicated. Genes shown represent the union of genes significantly regulated in mRNA-seq and/or Ser5-P CUTAC in at least one treatment comparison. mRNA-seq regulated genes were defined using adjusted P ≤ 0.001 and absolute log2FC ≥ 1; CUTAC regulated genes were defined using adjusted P < 0.05 and absolute log2FC ≥ 0.5. Genes were classified as RNA-only or CUTAC-only when significant regulation was detected in only one data type. Genes significantly regulated in both data sets were classified as concordant when RNA-seq and CUTAC changes occurred in the same direction, and discordant when they occurred in opposite directions. **D)** fGSEA of gene expression changes for protein-coding genes of (A). Top 15 gene sets across the indicated treatments are shown. Bubble color indicates normalized enrichment score (NES), bubble size indicates leading-edge fraction, and transparent bubbles indicate adjusted P ≥ 0.05. **E)** fGSEA of genes ranked by the non-additive transcriptional residual, defined as log2FC(FHT-1015/THZ531) − [log2FC(FHT-1015) + log2FC(THZ531)]. Gene sets with adjusted P < 0.05 are shown. Bubble color indicates normalized enrichment score (NES), and bubble size indicates leading-edge fraction. **F)** Gene-length distribution of additively and non-additively down-regulated (DOWN) and up-regulated (UP) genes of MDA-MB-231 cells co-treated for 8 h with FHT-1015 (750 nM) and THZ531 (300 nM) (n = 3). Genes were stratified using the non-additive transcriptional residual defined in (E), with |residual| < 0.5 classified as additive-like and |residual| ≥ 0.5 classified as non-additive. Boxplots show median and interquartile range; P values indicate Wilcoxon rank-sum tests.

Consistent with previous reports that CDK12 inhibition stimulates Pol II pause release and impairs Pol II processivity toward the 3′ ends of long genes,^10,12,67^ THZ531-treated cells showed a pronounced gene-length bias; up-regulated genes were significantly shorter and down-regulated genes longer than non-regulated genes. While BAF inhibition alone showed a similar but weaker trend, the combined CDK12 and BAF inhibition further exacerbated this bias for both up- and down-regulated genes (Fig. 5B). Likewise, exon-usage analysis confirmed this 5′-3′ bias in which 5′ exon up-regulation declined toward gene ends and 3′ exon down-regulation extended from the last two exon bins upon CDK12 inhibition to the last three upon combined CDK12–BAF inhibition (Fig. S5B). Together, these analyses show that BAF perturbation exacerbates the CDK12 inhibition-linked transcription defects, amplifying short gene induction and long gene repression.

We next asked whether transcription-coupled DNA accessibility changes corresponded to steady-state mRNA output. Clustering genes by their mRNA-seq and Ser5-P CUTAC responses to single-agent and combined CDK12–BAF inhibition revealed a larger group of concordantly regulated and a smaller group of discordant genes (Fig. 5C). Many genes showed directionally-matched mRNA-seq and Ser5-P CUTAC changes, but changes in one modality were often stronger, or statistical significant, without a matching significance change in the other. Thus, Pol II-coupled DNA accessibility and steady-state mRNA abundance are broadly linked in direction for many genes, but do not scale one-to-one in magnitude or statistical strength. We also asked whether BAF occupancy at promoters predicted transcriptional responses to CDK12 and BAF perturbation. While the induced genes were enriched for BRG1-occupied promoters, the down-regulated genes showed lower or non-significant enrichment (Fig. S5C). Thus, BAF presence at promoters marks accessible chromatin poised to gene induction but does not predict repression and may buffer against expression loss at selected loci.

To identify gene expression programs altered upon CDK12-BAF co-targeting, we performed gene set enrichment analysis (GSEA) on mRNA-seq log2 fold-changes from treated MDA-MB-231 cells using the Hallmark gene set collection of the Molecular Signatures Database.^68,69^ This analysis confirmed that the transcriptional response to combined CDK12 and BAF inhibition was largely defined by the effects of CDK12 inhibition alone (Fig. 5D). The strongest gene set enrichments were predominantly positive and included NF-κB, MYC, metabolic, epithelial mesenchymal transition, and adaptive-response gene programs.

We reasoned that the synergistic loss of cell viability upon combined CDK12 and BAF inhibition might involve transcriptional responses not captured by either single-agent treatment alone. To explore this, we identified gene-expression programs whose regulation in the combination deviated most strongly from the expected additive response by calculating an additive residual for each gene, defined as the observed log2FC after combined treatment minus the sum of the log2FC values observed upon the two single-agent treatments. Hallmark GSEA of the residual-ranked gene list revealed that NF-κB-dependent program, positively enriched in the combination relative to DMSO, was negatively enriched in the non-additive residual response (Fig. 5E). Thus, NF-κB signaling is induced by combined CDK12 and BAF inhibition less strongly than expected from the summed effects of the two single-agent treatments, suggesting that BAF inhibition partially constrains, rather than amplifies, the NF-κB-dependent transcriptional response upon CDK12 inhibition. Supporting this premise, chromatin proteomic analysis showed that BAF inhibition reduced levels of multiple signal-responsive TFs, including AP-1 family members JUN, JUNB, JUND, FOS, FOSB, FOSL1, FOSL2, ATF2 and ATF3, with concomitant increase of some AP-1 factors in the nucleoplasm (Fig. S5D). Since AP-1 and NF-κB cooperate in stimulating transcriptional programs,^70,71^ these data raise the possibility that reduced AP-1 chromatin engagement contributes to the constrained NF-κB residual response observed upon CDK12–BAF co-inhibition.

Importantly, the G2/M checkpoint and mitotic spindle programs were among the most strongly repressed non-additive responses to CDK12-BAF co-inhibition (Fig. 5E), suggesting that their down-regulation contributes to reduced viability of CDK12–BAF co-inhibited MDA-MB-231 cells. Notably, the non-additively regulated genes showed an even sharper gene-length bias than additively regulated genes: down-regulated genes were longer, whereas up-regulated were shorter (Fig. 5F).

Together, these data show that BAF ATPase inhibition modifies, rather than simply amplifies, the transcriptional response to CDK12 inhibition. The combined treatment retains a CDK12-dominated expression signature but adds a non-additive, gene-length-biased component characterized by enhanced induction of shorter genes and repression of longer genes involved in proliferation-associated programs.

### Induction of MYC mediates synthetic lethality upon combined CDK12 and BAF inhibition

Given that selective CDK12 inhibition renders cancer cells dependent on elevated Pol II pause release at oncogenic, signal-responsive gene programs,^67^ we reasoned that gene induction contributes to the loss of cell viability upon CDK12–BAF co-inhibition. Among the top short genes induced in MDA-MB-231 cells upon co-inhibition, oncogenic *MYC* caught our attention (Fig S6A). As a frequently amplified or overexpressed TF in several cancer types, including breast cancer, MYC is associated with tumor progression, relapse, and poor clinical outcome.^16,17^ In addition to controlling expression of distinct subsets of target genes,^72,73^ deregulated MYC also drives hypertranscription,^74^ collectively promoting proliferation, metabolic reprogramming, and other tumor-supportive gene expression programs. At the *MYC* locus, combined CDK12 and BAF inhibition increased transcription-coupled DNA accessibility, as shown by the elevated Ser5-P CUTAC signal at two hours of treatment (Fig. 6A), which persisted at later time points (Fig S6B). Correspondingly, mRNA-seq revealed synergistic induction of *MYC* mRNA upon combined CDK12 and BAF inhibition (Fig. 6B), accompanied by strong accumulation of MYC protein after 24 h in both whole-cell extract (Fig. 6C) and chromatin fractions (Fig. S6C). Induction of MYC, which occurred as early as four hours of combined treatment (*not shown*), did not translate into a proportional response of Hallmark MYC target genes, which were already strongly altered by CDK12 inhibition alone, which imposed the characteristic gene length-biased response, accentuated further by the combination (Fig. S6D).

**Figure 6.**
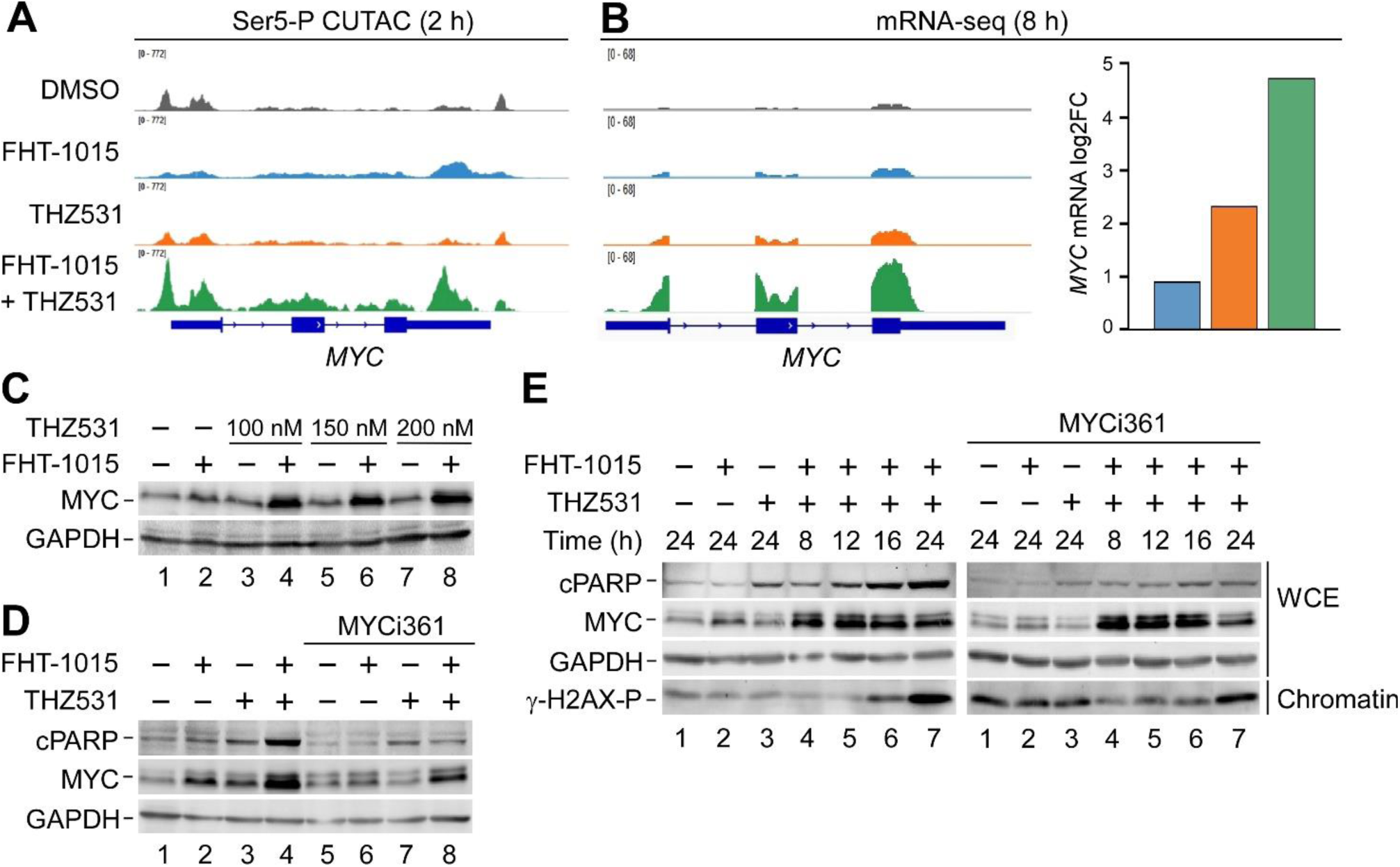
MYC induction contributes to apoptotic commitment upon combined CDK12 and BAF inhibition. **A)** Ser5-P CUTAC genome browser tracks at the *MYC* locus of MDA-MB-231 cells treated for 2 h with DMSO, FHT-1015 (1 μM), THZ531 (400 nM), and the combination as indicated. Each track shows a representative Ser5-P CUTAC replicate (n = 2). **B)** mRNA-seq genome browser tracks at the *MYC* locus (left) and *MYC* mRNA levels (right) of MDA-MB-231 cells treated for 8 h with DMSO, FHT-1015 (750 nM), THZ531 (300 nM), and the combination as indicated (n = 3). Each browser track shows a representative mRNA-seq replicate. DESeq2-derived MYC mRNA levels are plotted as the mean and relative to DMSO (adjusted P ≤ 0.001). **C)** Immunoblot analysis of MYC and GAPDH in whole cell extracts of MDA-MB-231 cells treated for 24 h with DMSO (–), FHT-1015 (750 nM), increasing concentrations of THZ531, and the combinations as indicated. **D)** Immunoblot analysis of the indicated proteins in whole cell extracts of MDA-MB-231 cells treated for 24 h with DMSO (–), FHT-1015 (750 nM), THZ531 (150 nM), and the combination with or without MYCi361 (5 μM) as indicated. **E)** Time-course immunoblot analysis of the indicated proteins in whole-cell extracts (WCE) and chromatin fractions of MDA-MB-231 cells treated with DMSO (–), FHT-1015 (750 nM), THZ531 (150 nM), and the combinations for the indicated durations with or without MYCi361 (5 μM) as indicated.

Aberrant MYC activity can also prime or activate cell death in a context-dependent manner, acting directly or indirectly on the mitochondrial and death-receptor apoptotic pathways.^75^ Analysis of pro-apoptotic and pro-survival gene expression changes in our mRNA-seq data revealed that the combination treatment significantly altered both gene classes but did not produce a simple pro-apoptotic transcriptional switch. Several stress-associated and pro-apoptotic genes, including *GADD45A*, *GADD45B*, *PMAIP1*/*NOXA*, *BBC3*/*PUMA,* and *BMF* were induced, whereas multiple apoptotic effector and death-receptor pathway genes, including *CASP3*, *CASP7*, *CASP8*, *TNFRSF10A/B*, and *TNFSF10*, were repressed (Fig. S6E). Among pro-survival genes, *XIAP*, *BCL2L1*, and *NAIP* were reduced by the combination, although this pattern largely resembled gene expression changes in response to CDK12 inhibition alone.

As mitochondrial pro-apoptotic regulator NOXA was previously linked to MYC-dependent apoptosis priming^76,77^, we next asked whether MYC induced by the combined CDK12 and BAF inhibition facilitated apoptosis. We monitored cPARP cleavage in whole-cell extracts of MDA-MB-231 cells treated with CDK12 and BAF inhibitors alone or in combination, with or without two orthogonal inhibitors that compromise MYC transcriptional activity. First, we used MYCi361, which disrupts MYC–MAX heterodimerization.^78^ Indeed, MYCi361 attenuated PARP cleavage induced by the combined treatment (Fig. 6D). This effect was recapitulated by CB-6644, a selective inhibitor of the ATPase activity of the MYC co-activator RUVBL1/2^79^ (Fig. S6F).

We next performed a 24 h time-course analysis to determine how MYC inhibition affected the kinetics of MYC accumulation, PARP cleavage, and γ-H2AX-P induction. As expected, PARP cleavage induced progressively by the combined CDK12 and BAF inhibition was blocked by MYCi361. Interestingly, MYC inhibition also increased γ-H2AX-P levels on its own and in combination with CDK12 and BAF inhibitor alone, but not when combined with CDK12 and BAF co-inhibition (Fig. 6E), suggesting that maintaining MYC activity above a critical threshold is important for preserving chromatin homeostasis in MDA-MB-231 cells. Together, these data show a key role of MYC in mediating the apoptotic response to combined CDK12 and BAF inhibition. Furthermore, our data suggest that cell death by apoptosis reflects a broader transcriptional imbalance rather than a simple, MYC-driven perturbation of genes that leads to apoptosis.

### Combined CDK12 and BAF inhibition impairs DNA synthesis and mitotic entry

The proliferative G2/M checkpoint and mitotic spindle programs were hallmarks of the non-additive, repressive responses to combined CDK12 and BAF inhibition (Fig. 5E). Coupled with augmented induction of shorter genes, a fraction of which are under MYC control, the balance between growth-promoting transcriptional outputs and the replication/cell-cycle machinery needed to sustain them may be perturbed, contributing to loss of cancer cell viability. This possibility is supported by the known role of deregulated MYC in increasing S-phase pressure and transcription–replication conflicts^80^, whereas CDK12 limits MYC-induced replication stress and maintains RNA Pol II processivity at DNA replication genes.^10,81^ Further, BAF perturbation has also been linked to replication stress and impaired fork progression.^82–84^ We therefore asked whether combined CDK12 and BAF inhibition alters DNA synthesis and cell-cycle progression.

To test this, we first measured DNA content and nascent DNA synthesis by DAPI/EdU flow cytometry after 8 h of inhibitor treatment, a time point that precedes detectable PARP cleavage in co-inhibited MDA-MB-231 cells (Fig. 7A and Fig. 3C). CDK12 inhibition combined with either BAF ATPase inhibition or BAF degradation reduced the fraction of EdU-positive S-phase cells beyond the additive expectation of the single treatments, with increased G2/M accumulation and only modest changes in G1 (Fig. 7B). At the same time, both combinations reduced EdU signal intensity within S-phase cells in a synergistic manner (Fig. 7C). Therefore, combined CDK12 and BAF inhibition reduces the fraction and DNA synthesis of replicating cells.

**Figure 7.**
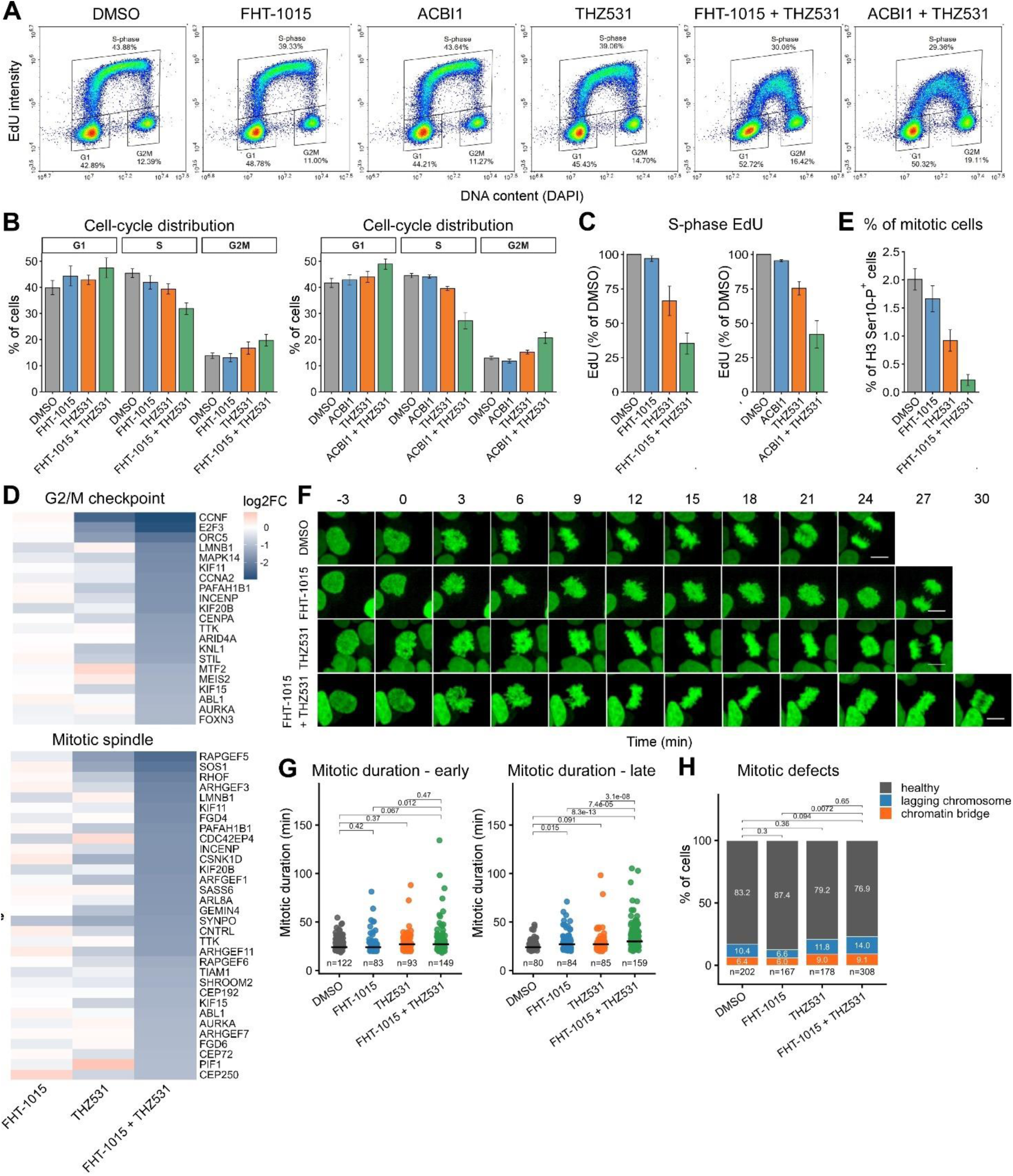
Combined CDK12 and BAF inhibition impairs DNA synthesis and mitotic entry. **A)** Representative EdU–DAPI flow cytometry plots after 8 h treatment with DMSO, 750 nM FHT-1015, 1000 nM ACBI1, 150 nM THZ531, or the indicated combinations. Gates indicate G1, S, and G2/M cell-cycle populations. FHT-1015/THZ531 dataset, n = 3; ACBI1/THZ531 dataset, n = 2. **B)** Quantification of cell-cycle distribution from DAPI DNA-content profiles of MDA-MB-231 cells treated for 8 h with DMSO, FHT-1015 (750 nM), ACBI1 (1 μM), THZ531 (300 nM), and the combinations as indicated. Left, FHT-1015:THZ531 dataset (n = 3); right, ACBI1:THZ531 dataset (n = 2). Data are plotted as the mean ± s.e.m. **C)** Quantification of EdU incorporation in S-phase MDA-MB-231 cells treated for 8 h with DMSO, FHT-1015 (750 nM), ACBI1 (1 μM), THZ531 (300 nM), and the combinations as indicated, shown as median EdU-A signal within the S-phase gate. Left, FHT-1015:THZ531 dataset (n = 3); right, ACBI1:THZ531 dataset (n = 2). Data are plotted as the mean ± s.e.m. **D)** Heatmaps showing mRNA-seq log2FC values (adjusted P ≤ 0.001) of non-additively downregulated G2/M checkpoint (top) and mitotic spindle genes (bottom) of MDA-MB-231 cells treated for 8 h with DMSO, FHT-1015 (750 nM), THZ531 (300 nM), and the combination as indicated, relative to DMSO (n = 3). **E)** Quantification of mitotic MDA-MB-231 cells treated for 8 h with DMSO, FHT-1015 (750 nM), THZ531 (300 nM), and the combination, measured as the percentage of phospho-histone H3 Ser10-positive cells by flow cytometry. Data are plotted as the mean ± s.e.m. (n = 3). **F)** Representative still images from live-cell imaging of MDA-MB-231 H2B-GFP cells treated with DMSO, FHT-1015 (750 nM), THZ531 (300 nM), and the combination, showing mitotic progression from nuclear envelope breakdown (NEBD) to anaphase onset. Time is shown in minutes relative to NEBD, which was set to 0 min. Scale bars, 10 μm. **G)** H2B-GFP imaging analysis of mitotic duration of MDA-MB-231 cells treated for 8 h with DMSO, FHT-1015 (750 nM), THZ531 (300 nM), and the combination as indicated. Mitotic duration was quantified for cells entering mitosis early (2-5 h) or late (5–8 h) after treatment. Each dot represents one mitotic event; median duration and event number are indicated for each condition. DMSO and FHT-1015 + THZ531 (n = 3); FHT-1015 and THZ531 (n = 2). P values were calculated using a two-sided Mann–Whitney test. **H)** Quantification of lagging chromosomes and chromatin bridges among the mitotic events scored in (G). P values were calculated using Fisher’s exact test comparing defective versus normal mitoses.

In line with previous work linking CDK12 to Pol II processivity at core G1/S genes,^10^ mRNA-seq analysis showed that CDK12 was required for expression of multiple DNA replication and S-phase progression factors (Fig. S7A-C). However, combined CDK12–BAF inhibition did not broadly exacerbate this gene-level repression, suggesting that the stronger DNA synthesis defect is unlikely to be explained by further loss of replication factors.

Importantly, combined CDK12–BAF inhibition repressed G2/M checkpoint and mitotic spindle genes in a synergistic manner (Fig. 7D and Fig. S7D). Among the genes repressed by the combination were regulators of cell-cycle progression and DNA replication-associated proliferation, including *E2F3, CCNA2, CCNF,* and *ORC5*, as well as genes required for mitotic entry and spindle function (*AURKA, TTK, KIF11, KIF15,* and *KIF20B*), chromosome segregation and kinetochore function (*INCENP, CENPA,* and *KNL1*), and centrosome duplication or organization (*SASS6, CEP192, CEP72,* and *CEP250*) (Fig. 7D). We validated *CCNA2* mRNA-seq data by immunoblotting, confirming that combined CDK12 and BAF inhibition reduces CDK1- and CDK2-activating Cyclin A2 levels in a synergistic manner (Figure S7E). We therefore asked whether CDK12 and BAF inhibition altered mitotic entry or progression. Flow-cytometric quantification of phospho-histone H3 Ser10, a marker of mitotic chromatin condensation, showed that that the combination reduced the fraction of mitotic cells by approximately ten-fold relative to untreated cells (Fig. 7E), indicating that the increased G2/M fraction (Fig. 7B) does not simply reflect accumulation of cells undergoing productive mitosis. This reduction is consistent with repression of genes required for G2/M progression and mitotic entry, particularly *CCNA2* and *AURKA*.^85^

To examine mitotic progression directly, we generated H2B-GFP-expressing MDA-MB-231 cells and used live-cell imaging to quantify mitotic duration and follow individual mitotic events. Mitoses were binned by mitotic entry time, defined by nuclear envelope breakdown, into early-entry and late-entry events occurring 2–5 h or 5–8 h after treatment, respectively. Among cells that entered mitosis, combined CDK12–BAF inhibition modestly prolonged mitotic duration, most clearly in the late-entry population (Fig. 7F,G). Overt mitotic defects, including lagging chromosomes and chromatin bridges, were modestly increased by the combination treatment (Fig. 7H). We conclude that, in addition to MYC-mediated apoptosis, impaired DNA synthesis and a strong block to mitotic entry are major consequences of CDK12–BAF co-inhibition, whereas the minority of cells that do enter mitosis show a modest delay without broad chromosome-segregation defects.

## DISCUSSION

Here, we identify BAF-dependent chromatin remodeling as a functional vulnerability in CDK12-inhibited TNBC cells. Genome-wide CRISPRi screening showed that CDK12 inhibition creates a dependency on canonical BAF subunits, while pharmacological co-targeting of CDK12 and BAF ATPase activity caused synergistic loss of viability across multiple TNBC models. Mechanistically, co-inhibition disrupted Pol II elongation and transcription-coupled chromatin accessibility. Further, while the co-inhibition exacerbated the CDK12-imposed, gene length-biased transcriptional response, it also modified it. The resulting transcriptional perturbation, marked by synergistic induction of short genes including MYC and repression of long cell cycle and mitotic genes, impaired DNA synthesis and mitotic entry, ultimately culminating in MYC-dependent apoptosis. Together, our findings identify the CDK12-BAF axis as a potential therapeutic vulnerability in transcriptionally addicted TNBC.

Selective nanomolar inhibitors of individual tCDKs have enabled kinase-specific functions and therapeutic interactions to be interrogated with a precision not possible with earlier pan-kinase inhibitors. Our study contributes to ongoing efforts to define vulnerabilities created by CDK12 perturbation and exploit them therapeutically in transcriptionally dependent cancers. Early siRNA-mediated knockdown studies established the essential role of CDK12 in promoting HR-mediated DNA repair pathway, providing a rationale for targeting CDK12-deficient tumors with PARP inhibitors and DNA-damaging chemotherapeutics.^26–28^ Subsequently, mechanistic studies, CRISPR-based and pharmacological screens uncovered synergy between CDK12 inhibition and a variety of perturbations, including CDK9 inhibition in colon cancer-derived cells,^67^ androgen-receptor antagonism in castration-resistant prostate cancer,^86^ RUNX1/CBFβ inhibition in melanoma,^87^ and BCL-2/BCL-xL and EGFR inhibition in TNBC.^88,89^ Our genome-wide CRISPRi screen extends this landscape by identifying opposing dependencies on ATP-dependent chromatin remodelers: whereas depletion of canonical BAF subunits sensitized TNBC cells to CDK12 inhibition, perturbation of NuRD-associated factors conferred relative resistance. Because BAF and NuRD generally promote accessible and repressed chromatin states, respectively, these findings suggest that survival of CDK12-targeted cells depends on the balance between opposing chromatin-remodeling activities.

The functional cooperation between CDK12 and BAF converged biochemically on Pol II elongation dynamics and apoptotic commitment. A central consequence of CDK12–BAF co-inhibition was the early and synergistic loss of chromatin-associated CTD Ser2P-Ser5P elongation mark, accompanied by accumulation of unphosphorylated Pol II on chromatin. The loss of the elongation mark preceded PARP cleavage and was not rescued by pan-caspase inhibition, placing the Pol II elongation defect upstream of apoptotic execution. Strikingly, chromatin proteomics further revealed depletion of multiple components of the pre-mRNA 3′-end processing machinery, including CSTF and CPSF factors, following CDK12 inhibition, which was further enhanced by BAF inhibition. Given the tight coupling between 3′-end processing and Pol II termination, ^61,62^ these changes raise the possibility that CDK12–BAF co-inhibition disrupts productive coupling of elongation-deficient Pol II to 3′-end processing and termination, thereby promoting persistence of transcriptionally dormant Pol II on chromatin. This model does not necessarily conflict with the reported induction of premature cleavage and polyadenylation following CDK12 inhibition,^19,20^ which likely reflects increased utilization of internal polyadenylation sites as a consequence of altered Pol II elongation rate. Rather, our findings suggest that BAF inhibition may perturb further the normal coupling between Pol II elongation, 3′-end processing and termination, potentially shifting the consequences of CDK12 inhibition toward persistence of transcription complexes on chromatin.

Consistent with the identified co-dependency between CDK12 and the BAF, our DepMap analysis showed that BAF-mutated cancer models are preferentially dependent on selected Pol II elongation and processivity factors, including SPT5, PAF1C subunits, and ELL1. These dependencies are mechanistically coherent with prior work placing BAF complexes in direct proximity to the elongation machinery. In yeast, Swi/Snf travels with elongating Pol II through coding regions and supports histone eviction during transcriptional elongation.^90^ In mammalian neurons, activity-assembled nBAF interacts with Pol II elongation complex, and its ATPase activity is required for productive elongation at immediate-early genes.^91^ Lastly, SPT5, one of the strongest DepMap dependencies in BAF-mutated models, has been shown to promote Pol II processivity along gene bodies and maintain enhancer architecture through physical interaction with BAF subunits.^63^ Thus, our findings extend BAF function beyond the promoter-proximal +1 nucleosome, into productive Pol II elongation. We propose that BAF activity helps Pol II navigate nucleosomal barriers during elongation, a requirement that becomes essential when CDK12-dependent elongation control is compromised.

Our chromatin profiling data suggest that the CDK12–BAF interaction is shaped by regulatory-element context. BAF inhibition broadly displaced BRG1 from both promoters and distal enhancers, whereas CDK12 inhibition caused a more selective and gradual reduction of BRG1 occupancy at promoters. Whether this reflects a direct effect of CDK12 activity or a secondary consequence of altered Pol II dynamics remains unresolved. At the level of transcription-coupled DNA accessibility, BAF perturbation was most evident at distal enhancers, where BAF ATPase inhibition reduced both BRG1 occupancy and Ser5-P CUTAC signal. However, BRG1 occupancy at promoters was only transiently reduced upon combined CDK12-BAF perturbation, whereas Ser5-P CUTAC signal shortly increased before returning to baseline. This seeming uncoupling may reflect active compensation by promoter-associated chromatin regulators, such as EP400/TIP60 which can restore promoter accessibility after BAF inhibition,^41^ or PHIP, which protects promoters from NuRD-mediated repression by promoting ubiquitination and removal of NuRD from chromatin upon loss of BAF activity.^52^ Thus, BAF inhibition strongly compromises enhancer-associated Pol II accessibility, whereas promoter-associated accessibility remains comparatively resilient under combined treatment.

The transcriptome-level changes were gene length-biased, with preferential induction of short genes and repression of long genes, consistent with BAF loss exacerbating a CDK12-linked elongation/processivity defect. *MYC*, a prominent short gene synergistically induced following CDK12-BAF co-inhibition, was required for apoptotic activation. *MYC* induction did not, however, produce a proportional response of a hallmark set of MYC target genes, suggesting that MYC accumulates in a transcriptional landscape that does not fully support a productive MYC-driven oncogenic program. This notion is consistent with a recent observation that the chromatin remodeler CHD1 was required to maintain accessible chromatin enabling a transcriptional program associated with cancer progression in MYC overexpressing breast cancer cells^92^. Thus, CDK12–BAF co-inhibition imposes a two-pronged constraint on induced MYC: MYC accumulates, but the Pol II elongation and chromatin-remodeling capacity required to channel MYC activity into a growth-supportive transcriptional program is impaired, converting MYC accumulation into an apoptotic vulnerability.

On the other hand, genes involved in cell-cycle regulation, DNA synthesis, and proliferation were readily repressed by the combination, which was marked by cell-cycle perturbation, reduced DNA synthesis, strongly impaired mitotic entry, and delayed mitotic progression as early as 8 h after CDK12-BAF co-inhibition. Notably, G2/M checkpoint and mitotic spindle programs were down-regulated beyond the additive expectation of the two single treatments, whereas regulation of S-phase genes in the combination largely resembled CDK12 inhibition alone, consistent with previous reports linking CDK12 inhibition to replication-associated defects.^10,93^ Thus, the combination did not create a distinct S-phase transcriptional signature beyond CDK12 inhibition alone; rather, BAF inhibition exacerbated the functional consequences of CDK12-linked replication stress, consistent with a chromatin-level requirement for BAF in replication-fork progression and transcription–replication conflict control.^82–84^

More broadly, our findings support the therapeutic exploitation of Pol II elongation control as a cancer vulnerability shaped by chromatin state. Transcriptional elongation is increasingly recognized as a regulated and druggable step of gene expression, controlled by CDK9, CDK12/13, BRD4, SEC, SPT5, SPT6, PAF1C, Integrator, and associated chromatin regulators.^5^ In this framework, cancer cells may be sensitive not to inhibition of one transcriptional kinase or one remodeler, but to combinations that collapse compensatory mechanisms maintaining productive transcription through chromatin. Our data place BAF perturbation in this category by showing that disruption of ATP-dependent chromatin remodeling sensitizes TNBC cells to compromised CDK12-dependent elongation control. This suggests that chromatin-remodeling defects, whether pharmacologically induced or genetically encoded, could define cellular states with heightened dependency on elongation machinery. Conversely, selective targeting of elongation kinases or elongation-associated factors may create actionable vulnerabilities in cancers already carrying defects in chromatin remodeling. Thus, combined targeting of chromatin remodeling and transcription elongation may convert the transcriptional plasticity that normally supports cancer-cell adaptation into a therapeutic liability.

In conclusion, our study identifies BAF-dependent chromatin remodeling as an actionable vulnerability in CDK12-perturbed TNBC cells and links this functional interaction to impaired Pol II elongation, persistence of chromatin-associated Pol II, gene length-dependent transcriptional perturbation, defective DNA synthesis, and apoptotic commitment. These findings support a model in which BAF activity helps sustain productive transcription through chromatin when capacity to elongate is compromised, extending the role of BAF beyond promoter-proximal chromatin regulation into the maintenance of productive Pol II transcription. Future work should define which BAF subcomplexes and genomic contexts underlie this vulnerability, determine whether BAF-mutated tumors display heightened sensitivity to elongation-targeted therapies, and establish whether biomarkers of Pol II processivity, BAF status, or MYC-dependent stress can guide therapeutic co-targeting of chromatin remodeling and transcription elongation in cancer.

## MATERIALS AND METHODS

### Cell lines

MDA-MB-231, MDA-MB-436, HCC1395 breast cancer cell lines and HEK293T cells were cultured in RPMI-1640 (Corning, cat. no. 10-040-CV) and DMEM (Merck) medium, respectively, supplemented with 10% fetal bovine serum (FBS) and 100 U/ml penicillin/streptomycin. All cells were maintained at 37°C in a humidified incubator with 5% CO₂. Cell lines were regularly confirmed to be mycoplasma-free using a published PCR-based detection protocol.^94^ The MDA-MB-231 cell line was authenticated by short tandem repeat (STR) profiling by Eurofins. MDA-MB-436 and HCC1395 cells were not authenticated in-house but were obtained from trusted sources, as listed in Supplementary Table S1D.

### Chemicals

THZ531, FHT-1015, ACBI1, MYCi361 and CB-6644 were purchased from MedChemExpress. SR-4835 was a gift from Matthias Geyer Laboratory (University of Bonn, Germany). For more information on the chemicals, see Supplementary Table S1B.

### Lentivirus production and generation of stable cell lines

To prepare lentivirus for CRISPRi screen, HEK293T cells were transfected in 15 cm dishes using JetPrime transfection reagent (Polyplus), together with the packaging plasmid psPAX2 and the envelope plasmid pMD2.G. Lentivirus for CRISPRi validation experiments and stable cell line generation was produced using PEI MAX 40K (Polysciences #24765). Viral supernatants were used directly for transduction without concentration or polybrene. The MDA-MB-231 CRISPRi cell lines was generated by transduction with a lentivirus encoding ZIM3-KRAB-dCas9-(Addgene #154472), followed by selection with 5 µg/mL blasticidin (ThermoFisher) for 7 days and FACS sorting to obtain single-cell clones. For CRISPRi validation experiments, individual sgRNAs oligonucleotides (Integrated DNA Technologies) were subcloned into a lentiviral sgRNA expression vector, and transduced cells were selected for 3 days with 2 µg/mL puromycin (ThermoFisher). For live-cell imaging, an MDA-MB-231 H2B-GFP-expressing population was generated by transduction with lentivirus encoding PGK-H2B-eGFP (Addgene #21210) and used as a polyclonal population without antibiotic selection.

### CRISPRi screening

Lentivirus of the pooled CRISPRi Dolcetto sgRNA library (Addgene Cat# 92385 and Cat# 92386) was produced as described above. 90 million dCas9-ZIM3-KRAB MDA-MB-231 cells were transduced in cell culture media at MOI of 0.3. Cells were expanded for two days and selected with 2 µg/mL puromycin for three days. Cells were cultured in the presence of IC50 (110 nM) THZ531, or corresponding DMSO volume, for 18 days. Library representation of 500x was maintained. Cells at day 0 and day 18 were collected and frozen −80°C until further processing. DNA was extracted with the NucleoSpin Blood L Midi kit (Macherey-Nagel) and quantified on a Nanodrop Spectrophotometer (Thermo Fisher Scientific). All purified DNA was amplified with NEBNext® Ultra™ II Q5 polymerase in 100µl reactions, making sure DNA per reaction does not exceed 20 µg. PCR cycles were 1× (98 °C, 30 sec), 24× (95 °C, 15 s; 65 °C, 20 s; 72 °C, 20 s), 1× (72 °C, 1 min). Forward primers are an equimolar mix of eight staggered primers (5′-AATGATACGGCGACCACCGAGATCTACACTCTTTCCCTACACGACGCTCTTCCGATCT-(N0-8)-TTGTGGAAAGGACGAAACACCG-3′), reverse primers include an eight-nucleotide long sample specific index (5’-CAAGCAGAAGACGGCATACGAGAT-(N8)-TTGTCCGAGTGACTGGAGTTCAGACGTG-3’) and both are used at a 0.25 µM final concentration. PCR reactions are cleaned up using the Monarch Spin PCR & DNA Cleanup Kit (NEB). Purified PCR products were run on 1.5% agarose gel, followed by gel extraction and another round of PCR cleanup purification and quantified with the Qubit dsDNA HS Assay Kit. DNA was normalized and pooled. Pooled DNA was diluted to a final concentration of 10 nM. Libraries were sequenced at the CZ Biohub San Francisco on a NovaSeq 6000 (Illumina) with read 1 and read 2 lengths of 70 and 30 bases respectively.

### Analysis of CRISPR screen

Read preprocessing and counting of the sequencing data was performed as follows. Forward reads were first trimmed with BBDuk (v. 38.41, parameters: ktrim=l minlen=20 k=10 mink=6 hdist=1 tpe tbo skipr2) to remove the human U6 sequence trailing the spacer sequence (5’-TTGTGGAAAGGACGAAACACCG-3’). Forward and reverse reads were hard clipped to 20 and 21 bases respectively. Forward reads were counted for perfect matches to the guide RNA library using R (v. 4.5.0) with the packages ShortRead (v. 1.67.0)^95^ and dplyr (v. 1.1.4). Gene-level effect sizes were estimated with MAGeCK Robust Ranked Aggregation (v. 0.5.7)^50^ using non-target controls as control sgRNA.

### Validation of CRISPRi screens using individual gRNA competition assay

To independently validate CRISPRi screen hits, sgRNAs targeting genes of interest were individually cloned into an sgRNA expression vector by Gibson assembly. A full list of sgRNA sequences is provided in Supplementary Table X. To assay differential sensitivity to THZ531, test sgRNAs were cloned into an mCherry-expressing sgLenti vector, while a non-targeting control sgRNA was cloned into a GFP-expressing sgLenti vector.^96^ Cells were lentivirally transduced and selected with puromycin. mCherry-positive cells expressing the test sgRNA were mixed with GFP-positive control cells at a defined ratio: 4:1 for BAF-targeting sgRNAs and 1:1 for NuRD/NuRD-associated sgRNAs. Mixed cell populations were treated with THZ531 at the IC50 concentration, 110 nM, or DMSO-treated, and the ratio of mCherry-to GFP-positive cells was monitored by flow cytometry over two weeks. BAF-targeting sgRNA competition assays were performed from two independent lentiviral transductions, each measured in three technical replicate wells, and plotted as mean ± s.e.m. NuRD/NuRD-associated sgRNA competition assays were performed from one lentiviral transduction measured in three technical replicate wells, and plotted as mean ± s.d.

### Cytotoxicity and viability assays

Cytotoxicity and viability were measured using the CellTox Green Cytotoxicity Assay (Promega) and CellTiter-Glo 2.0 Cell Viability Assay (Promega), respectively, according to the manufacturer’s instructions. Cells were seeded in 96-well plates at a density of 15,000 cells per well 16 h before treatment. Fluorescence and luminescence were measured at the indicated time points using a PerkinElmer Victor X3 plate reader.

For viability measurements, CellTiter-Glo values were normalized to DMSO-treated cells, defined as 100% viability, and cells treated with 40 µM benzethonium chloride, defined as 0% viability. Percent viability was calculated as:

% viability = 100 − [((signal − DMSO signal)/(benzethonium chloride signal − DMSO signal)) × 100].

For cytotoxicity measurements, CellTox Green values were normalized to DMSO-treated cells, defined as 0% cytotoxicity, and cells treated with 40 µM benzethonium chloride, defined as 100% cytotoxicity. Percent cytotoxicity was calculated as:

% cytotoxicity = ((signal − DMSO signal)/(benzethonium chloride signal − DMSO signal)) × 100. Drug synergy was calculated using SynergyFinder^55^ based on the Bliss independence model. Presented values represent the mean three independent biological experiments.

### Cell confluency assay

The assay was performed using the CELLCYTE X™ live cell imager and analyzer (Cytena). All cell lines were seeded in 96-well plates at a density of 3,000 cells per well for 16 h prior to the treatment. Cells were imaged every four hours for seven days. Cell culture medium and drugs were replenished on day 3 of the assay. Confluency values were quantified using the CELLCYTE X™ software. Cell confluency assays show one representative experiment measured in three technical replicate wells and are plotted as mean ± s.e.m.

### Organoid viability assay

Following previously described methods,^57^ TORG40 organoids were grown in 50 μL 100% BME gel domes in 24-well plates supplemented with organoid medium and grown to full confluency. One confluent well in a 24-well plate was used to make 10–20 wells in a 96-well plate. Organoids were collected and digested into small clumps using TrypLE for 15 min, filtered through a 100 μm filter, and resuspended in organoid medium containing 6% BME. 90 μL of organoid suspension was dispensed into each well of a 96-well plate, with the outermost wells left unused. On day 2, 30 μl of medium containing THZ-531 and/or FHD-286 (single and combination doses) in DMSO was added, in at least technical triplicate. The final DMSO concentration in the wells was kept under 1%. Drug experiments were stopped at 72 h, and cell viability was assessed using CellTiter-Glo.

### RNA extraction and mRNA-sequencing

Cells were seeded on 6-well plates for 16 h and treated at 80% confluency with drug doses that provided the most synergistic effect on cell viability and cytotoxicity - 750 nM FHT-1015, 300 nM THZ531 – alone or in combination, alongside the corresponding volume of DMSO. The cells were harvested 8h post-treatment – a time point at which there was no decline in cell viability, as measured by CellTiter-Glo 2.0 Cell Viability Assay (Promega). RNA samples were extracted using RNeasy Mini Kit (Qiagen) and treated with RNase-Free DNase (Qiagen). The extracted RNA was quality controlled with Agilent 2100 Bioanalyzer, showing characteristic total RNA profiles and RIN values between 9.20 and 10.00. Subsequent selection of polyadenylated RNAs and library preparation was conducted by National Genomics Infrastructure (NGI) Sweden, using Illumina’s TruSeq polyA selection kit. The mRNA-seq libraries were sequenced with NovaSeq XPlus using paired-end (2×150 bp) layout. RNA-seq samples were processed using the nf-core/rnaseq pipeline.^97^ Quality control and adapter trimming was peformed using Trim Galore (https://www.bioinformatics.babraham.ac.uk/projects/trim_galore/) and reads were aligned against the hg38 human genome using STAR.^98^ Number of read counts per gene and exon were determined from the mapped RNA-seq reads in a strand-specific manner using featureCounts^99^ and gene annotations from Ensembl version 100.^100^ Differential gene expression analysis was performed using DESeq2.^101^ Differential exon usage was determined using.^102^ P-values were adjusted for multiple testing using the method by Benjamini and Hochberg.^103^ Genes were classified as significantly regulated using adjusted P ≤ 0.001 and absolute log2FC ≥ 1. Exons with an adjusted P ≤ 0.01 were classified differentially used. Differential analysis was performed with a Watchdog workflow.^104^ Gene set enrichment analysis was performed using the fgsea R package^105^ with Hallmark gene sets from the Molecular Signatures Database.^68^ Genes were ranked by RNA-seq log2FC values or by the non-additive transcriptional residual, as indicated. Gene sets with adjusted P < 0.05 were considered significant. For treatment-versus-DMSO analyses, genes were ranked by RNA-seq log2FC values. For analysis of the non-additive transcriptional response, genes were ranked by the transcriptional residual calculated as the observed log2FC after combined FHT-1015/THZ531 treatment minus the sum of log2FC values after the two single-agent treatments. Gene sets with adjusted P < 0.05 were considered significant.

### CUT&Tag and Ser5-P CUTAC chromatin profiling

CUT&Tag and Ser5-P CUTAC were performed using a CUT&Tag-direct whole-cell protocol with minor modifications.^105,106^ Cells were seeded on 6-well plates for 16h and treated at 60% confluency with 400 nM THZ531, 1 uM FHT1015, or combination, for the indicated duration. Untreated cells collected at 0 h were used as the reference condition. Cells were detached by trypsin and inactivated using FBS-containing medium. The cell suspension was transferred to a 15 mL tube and centrifuged at 600xg for 3 minutes. Supernatant was removed and the cells were resuspended in 2 mL 1X PBS and counted using the ViCell Cell Counter.

For each reaction, 50,000 cells with viability greater than 90% were bound to 20 µL concanavalin A magnetic beads for 10 min. Cell-bound beads were incubated overnight at 4°C with primary antibody diluted 1:100 in Antibody Buffer (20 mM HEPES pH 7.5, 150 mM NaCl, 0.5 mM spermidine, 0.05% Triton X-100, 0.1%BSA, 2 mM EDTA). Antibodies against BRG1/SMARCA4 (Abcam) and CDK12 (Sigma-Andrich) were used for CUT&Tag, and an antibody against RNA polymerase II Ser5-phosphorylated CTD (CellSignaling) was used for CUTAC. After primary antibody incubation, cells were incubated with guinea pig anti-rabbit secondary antibody diluted 1:100 in Triton Wash Buffer (20 mM HEPES pH 7.5, 150 mM NaCl, 0.5 mM spermidine, 0.05% Triton-X100 and Roche EDTA-free protease inhibitor) for 1 h at room temperature. Cells were washed with Triton Wash Buffer and incubated with CUTANA pAG-Tn5 transposase (EpiCypher) diluted 1:20 in Triton 300-wash buffer (Triton Wash buffer with 300 mM NaCl) for 1 h at room temperature, followed by a wash with Triton 300-wash buffer.

For CUT&Tag, tagmentation was performed in Triton 300-wash buffer supplemented with 10 mM MgCl₂ for 1 h at 37°C. For CUTAC, tagmentation was performed in CUTAC-DMF tagmentation buffer (10 mM TAPS, 5 mM MgCl_2_, 20% DMF, 0.05% Triton-X100) for 20 min at 37°C. Tagmented DNA was released using SDS–proteinase K release solution, followed by neutralization with Triton X-100. Libraries were amplified using NEBNext High-Fidelity 2× PCR Master Mix and indexed Illumina adapters, purified using 1.3× SPRI bead cleanup, and resuspended in 10 mM Tris-HCl pH 8. Library size distribution was assessed using an Agilent 4200 TapeStation with D1000 reagents. Barcoded libraries were pooled and sequenced as paired-end 50 bp reads.

CUT&Tag and Ser5-P CUTAC data have been deposited to Gene Expression Omnibus (GSE311170).

### CUT&Tag and Ser5-P CUTAC data processing

Paired-end reads were adapter-trimmed using cutadapt and aligned to the hg19 human genome assembly using Bowtie2 with the--very-sensitive-local,--soft-clipped-unmapped-tlen, and --dovetail options.^107,108^ Genome tracks were generated as library-size-normalized bedGraph files and visualized using Integrative Genomics Viewer. BRG1 and CDK12 CUT&Tag peaks were called from replicate samples using SEACR^109^ with stringent peak calling and a top 5% threshold. Peaks were ranked by total signal, and the top 20,000 BRG1 and CDK12 peaks were used for downstream comparisons.

Peaks were annotated by overlap with ENCODE candidate cis-regulatory elements, excluding regions overlapping masked genomic regions.^110^ Promoters and distal enhancers were classified as BRG1-only, CDK12-only, or shared BRG1/CDK12-bound regulatory elements based on overlap between annotated BRG1 and CDK12 peak sets. Summary profiles and heatmaps of normalized CUT&Tag and CUTAC signal were generated using deepTools.^111^ For differential CUTAC analysis at genes and distal enhancers, normalized count matrices were generated and analysed using DESeq2.^101^ Genes and enhancers with no detectable counts were excluded. K-means clustering was used to group genes and enhancers according to CUTAC signal changes over time.

### Integrated mRNA-seq and Ser5-P CUTAC analysis

Gene-level RNA-seq and Ser5-P CUTAC log2FC values were integrated to compare steady-state transcriptional changes with transcription-coupled chromatin accessibility after 8 h treatment. Genes detected in both datasets were retained. RNA-seq-regulated genes were defined using adjusted P ≤ 0.001 and absolute log2FC ≥ 1, and S5-P CUTAC-regulated genes were defined using adjusted P < 0.05 and absolute log2FC ≥ 0.5. Genes were classified as RNA-seq-only, CUTAC-only, concordant, or discordant based on the significance and direction of regulation in the two datasets.

### Intracellular fractionation for mass spectrometry

After incubation with tested compounds for the indicated duration, MDA-MB-231 cells were washed and scraped with ice-cold PBS. Cell pellets were fractioned as previously described.^112,113^ The chromatin pellets were incubated with 2 uL Pierce™ Universal Nuclease (Thermo) for 1h at 4°C. Resulting nucleoplasm and chromatin fractions were precipitated using ProteoExtract precipitation kit (Millipore) according to manufacturer’s instructions and snapped-frozen until MS analysis.

### Mass spectrometry acquisition and analysis

Acetone precipitated protein samples were treated as reported.^114,115^ Briefly, cells were lysed in 5 % sodium dodecyl sulfate (SDS), reduced with 10mM Tris (2-carboxyethyl) phosphine hydrochloride (TCEP) and alkylated with 40mM chloroacetamide (CAA) in 100 mM triethylammonium bicarbonate (TEAB) containing protease and phosphatase inhibitors (Roche). Proteins were heated for 10 min at 95 °C and sonicated for 15 min with Bioruptor (Diagenode) using 30 s on/off cycles. Proteins were aggregated to MagReSyn hydroxyl beads (Resyn Biosciences) using acetonitrile. Proteins were digested overnight using trypsin/Lys-C mix (Promega). An aliquot of the sample was taken for total protein analysis and for remaining sample phosphopeptide enrichment was carried out using High Select Fe-NTA kit (Thermo Fisher Scientific) according to manufacturer’s instruction.

The LC-ESI-MS/MS analysis was performed on Evosep One HPLC system (Evosep, Odense, Denmark) coupled to the Orbitrap Astral mass spectrometer (Thermo Fisher Scientific, Bremen, Germany) equipped with a nano-electrospray ionization source. Peptides were separated inline on an Aurora Elite C18 UHPLC column (15 cm x 75 µm, Ionopticks). The mobile phase consisted of water with 0.1% formic acid (solvent A) and 0.1% formic acid/99.9% acetonitrile (*v*/*v*) (solvent B). Samples were analysed with 40 samples per day Whisper Zoom method using a data independent acquisition (DIA) LC-MS/MS method. MS data was acquired automatically by using Thermo Xcalibur software 4.7 (Thermo Scientific). In a DIA method, a duty cycle contained one full scan range of 400–900 *m/z* with resolution of 180 000. Normalized full-MS AGC target was set to 500%. Fragment ions were scanned with a range of 400.4319.428–900.6593 *m/z* using 142 DIA MS/MS scans with variable width isolation windows. MS2 fragmentation was achieved by normalized collision energy of 25% and fragment scans were recorded at maximum fill time 2.5 ms.

Data analysis consisted of protein identifications and label free quantifications of protein abundances. Data was analysed by Spectronaut software (Biognosys; version 20.1.250624.92449) against *Homo sapiens* database (SwissProt, release 2025_03) and Universal Protein Contaminant database.^116^ DirectDIA approach was used to identify proteins and label-free quantifications were performed with MaxLFQ. Trypsin/P was selected as the enzyme and maximum of two missed cleavages were allowed. Carbamidomethylation of cysteine was chosen as fixed modification and protein N-terminal acetylation and oxidation of methionine were selected as variable modifications. Precursor and protein FDR cutoff were set to 0.01. Quantification was carried out on MS2 level.

For total proteomics analysis, protein-group quantity tables were analysed in R. Protein intensities were normalized within each sample by total protein abundance and log2-transformed. Proteins were retained for differential analysis if they were quantified in at least two samples in any condition–fraction group. Chromatin-associated and nucleoplasmic fractions were analysed separately using limma. For each fraction, linear models were fitted to compare drug treated samples with the corresponding DMSO control. Differentially enriched or depleted proteins were ranked by log2 fold change and moderated statistics from limma, and results were visualized as volcano plots.

### Western blotting

Whole-cell extracts were prepared using lysis buffer (50 mM Tris-HCl, 0.5% NP-40, 150 mM NaCl, 1 mM EDTA, pH 7.4) on ice for 45 mins in the presence of Protease Inhibitor Cocktail (MedChemExpress). Lysates were cleared by centrifugation at 16,000 g for 15 min, boiled in SDS loading buffer supplied with 10% of β-Mercaptoethanol for 5 min, separated using SDS-PAGE, transferred to nitrocellulose membrane and analyzed by Western blotting. All primary antibodies were used at 1:1000 dilutions. Manufacturers provide validation for all antibodies. For more information on the antibodies, see Supplementary Table S1A.

### Obtaining chromatin fraction for western blotting

The nuclear fraction from drug- and DMSO-treated cells was obtained using the published REAP protocol^117^ and further processed as described by Devaiah and colleagues.^118^ The chromatin pellet was resuspended in 100 uL ice-cold buffer A (10 mM HEPES pH 7.9, 1.5 mM MgCl2, 10 mM KCl, 75 mM NaCl, 1 mM DTT, 0.5% NP-40, Protease Inhibitor and Phosphatase Inhibitor Cocktail) and incubated with 2 uL Nuclease for 1h at 4°C. Digested chromatin was boiled in SDS loading buffer.

### EdU incorporation, DNA-content and phospho-H3 flow cytometry

Cells were seeded on 6-well plates for 16 h and treated at 80% confluency with the indicated compounds and corresponding volume of DMSO for 8h. EdU was added for 30 min at a final concentration of 20 µM. Cells were harvested, fixed using Click-iT fixative and EdU incorporation was detected using the Click-iT EdU Alexa Fluor 488 Flow Cytometry Assay Kit (ThermoFisher) according to the manufacturer’s instructions. Cells were stained with anti-phospho-Histone H3 Ser10 antibody followed by CoraLite647-conjugated secondary antibody, and DNA content was stained with DAPI. Flow cytometry was performed using the Novocyte Quanteon cytometer and data were analysed using the Novocyte software. Cell-cycle distribution was quantified from DAPI DNA-content profiles, EdU incorporation was quantified as median EdU-A signal within the S-phase gate, and phospho-H3-positive cells were quantified from the Ser10 phospho-H3-positive population. Quantified data are presented as mean ± s.d. from the indicated number of biological replicates.

### Live-cell imaging

Time-lapse imaging was performed using an Opera Phenix Plus high-content screening system (Revvity) in confocal mode equipped with a 40× water immersion objective (NA 1.1). EGFP fluorescence was acquired using 488 nm excitation and a 500–550 nm emission filter set. Images were collected with a field of view of 2160 × 2160 pixels at binning 1. A total of 42 fields were imaged across 20 z-planes per field over 120 timepoints from cells cultured in an Ibidi µ-slide chamber.

Mitotic events were scored manually from time-lapse image sequences of MDA-MB-231 H2B-GFP cells. Mitotic duration was measured from nuclear envelope breakdown to anaphase onset. Mitotic phenotypes were classified as normal mitosis, lagging chromosomes, or chromatin bridges based on H2B-GFP chromosome morphology during mitosis.

### DepMap CRISPR dependency analysis

DepMap model metadata, CRISPR gene-effect scores, and somatic mutation annotations were imported from DepMap. Only cancer models were retained for analysis. Mutation status was defined using variants annotated as HIGH impact by VEP. For the pan-BAF analysis, models were classified as BAF-mutated if they carried a HIGH-impact mutation in any of the following canonical BAF-related genes: SMARCA4, SMARCA2, SMARCB1, SMARCC1, SMARCC2, SMARCD1, SMARCD2, SMARCD3, SMARCE1, ACTL6A, ACTL6B, ACTB, ACTG1, ARID1A, ARID1B, DPF1, DPF2, DPF3, or PHF10. All remaining cancer models without HIGH-impact mutations in this gene set were classified as BAF-WT/non-mutated.

For subgroup analyses, mutation groups were defined using HIGH-impact mutations in selected individual BAF subunits or paralogous subunit groups. The displayed subgroup analyses include models with HIGH-impact mutations in SMARCC1 or SMARCC2, and models with HIGH-impact mutations in ARID1A or ARID1B. These groups were compared with the corresponding non-mutated cancer models.

CRISPR gene-effect scores for selected Pol II transcription-cycle regulators were compared between mutated and non-mutated groups. Differential dependency was quantified as the difference in median CRISPR gene-effect score between the two groups. Negative Δ median values indicate stronger dependency in the mutated group. Statistical significance was assessed using two-sided Wilcoxon rank-sum tests. Because the DepMap analysis was used as an exploratory survey, nominal P values are shown without multiple-testing correction. Results were visualized as ranked dependency plots showing Δ median CRISPR gene effect, with point size representing −log10(nominal P).

### Quantification and statistical analysis

Unless otherwise stated, statistical analyses were performed in R^119^, and data were visualized using ggplot2^120^. The number of biological replicates, technical replicate wells, or scored events is indicated in the corresponding figure legends. Data are presented as mean ± s.e.m., mean ± s.d., individual values, or medians as indicated in the figure legends. Gene-length distributions were compared using Wilcoxon rank-sum tests. Mitotic duration was compared using a two-sided Mann–Whitney test. Mitotic defect frequencies were compared using Fisher’s exact test. DepMap dependency comparisons were performed using two-sided Wilcoxon rank-sum tests. For exploratory DepMap analyses, nominal P values are shown without multiple-testing correction. Method-specific statistical thresholds for CRISPRi screening, RNA-seq, CUTAC, proteomics, drug synergy, and gene set enrichment analyses are described in the corresponding methods sections.

## CONFLICT OF INTEREST DISCLOSURE

The authors declare no conflict of interest.

## ACKNOWLEDGEMENTS

We thank the McManus laboratory for hosting M.M. and providing a supportive research environment during the CRISPRi screening work; M.Geyer for sharing SR-4385; Members of Barboric laboratory for support throughout the project; FIMM High Content Imaging and Analysis Unit (FIMM-HCA), HiLIFE, University of Helsinki, Euro-Bioimaging and Biocenter Finland for imaging services; Mass spectrometry analysis by Turku Proteomics Facility, University of Turku and Åbo Akademi University, supported by Biocenter Finland; and all the authors for critical reading of the manuscript.

## AUTHOR CONTRIBUTIONS

M.B. and M.M. conceptualized the study and designed the experiments. M.B. supervised the study. M.M. performed and analyzed the CRISPRi screen with the assistance of S.O. under the supervision of M.T.M. M.M. performed genetic and pharmacological cell growth, viability, and cytotoxicity assays across TNBC model systems. M.M. conducted immunoblotting analyses, cell-cycle progression, DNA synthesis, and live-cell imaging experiments, together with downstream quantification. H.Y and Y.B performed 3’-end processing factors and Cyclin A immunoblotting analyses, respectively. T.B.V.B performed and analyzed organoid experiments under the supervision of J.M.R. C.D. performed and analyzed the CUT&Tag and Ser5-P CUTAC under the supervision of S.H. M.M. prepared samples for mRNA-seq and mass spectrometry analyses. O.K. processed the samples for mass spectrometry and performed data acquisition and primary data processing. C.C.F. performed gene-expression analyses of the RNA-seq data. M.M. and M.B. wrote the manuscript with assistance from M.T.M., S.H., C.C.F., A.V., and S.O.

## FUNDING

Research Council of Finland [1361495 to M.B.]; Sigrid Juselius Foundation [4707108 to M.B.]; Cancer Foundation Finland [4708690, 4706832, and 4707579 to M.B.]; Instrumentarium Foundation [to M.B.]; Magnus Ehrnrooth Foundation [4706635 and 4707773 to M.B.]; Rivkin Center for Ovarian Cancer [4721267 to M.B.]; Deutsche Forschungsgemeinschaft (FR 2938/14-1 and FR2938/11-2 in the framework of the Research Unit FOR5200 DEEP-DV (443644894)) [to C.C.F.]; California Breast Cancer Research Program [to J.M.R.]; National Institutes of Health grants [R01 CA279801 and U01 CA272546 to M.T.M]; Howard Hughes Medical Institute [to S.H.].

## SUPPLEMENTARY FIGURES AND FIGURE LEGENDS

**Figure S1.**
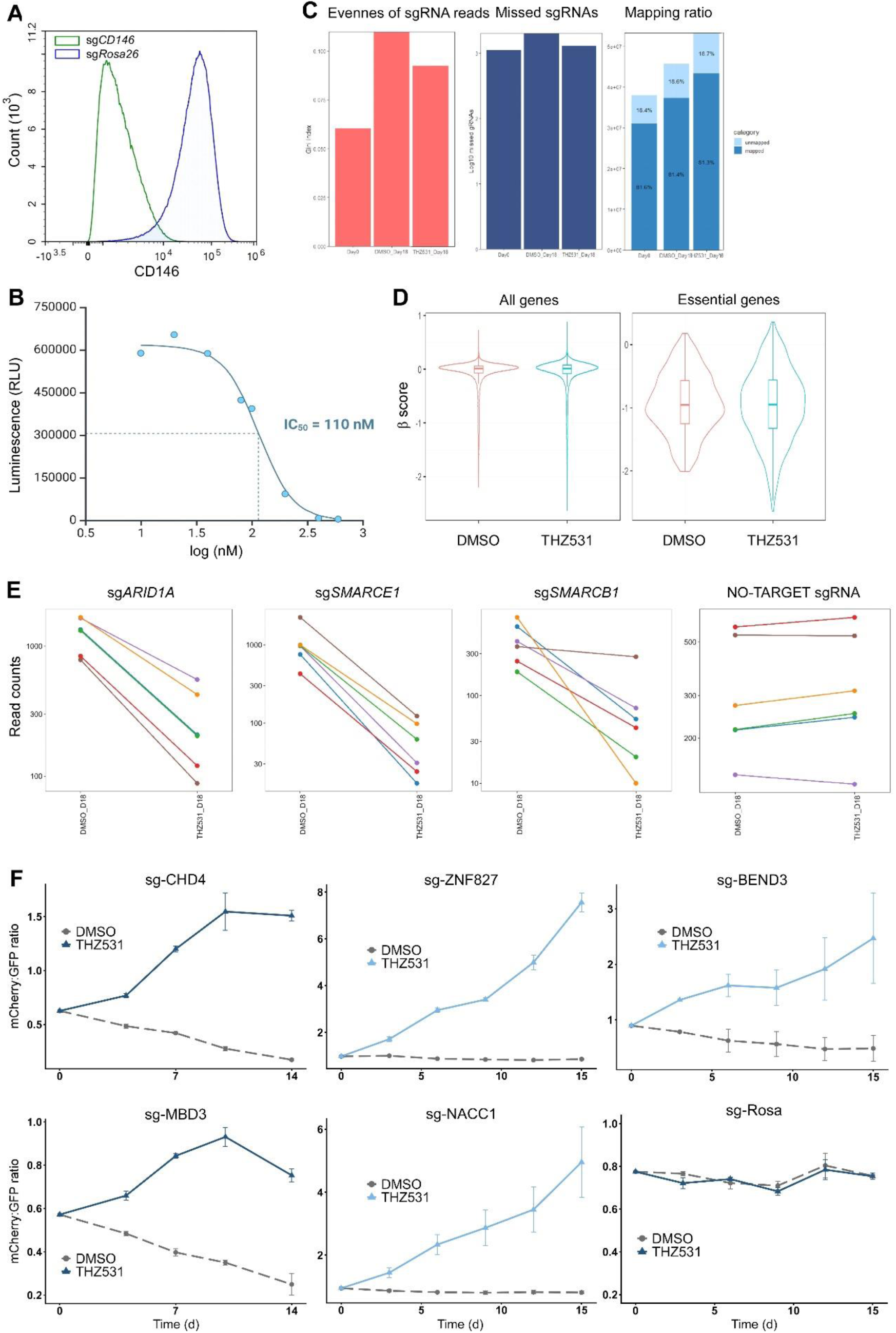
Quality control and validation of the genome-wide CRISPRi screen. **A)** Protein levels of cell-surface antigen CD146 in MDA-MB-231 CRISPRi monoclonal cell line transduced with sgRNA targeting *CD146* or *Rosa26* as determined by antibody staining and flow cytometry. **B)** THZ531 dose–response curve in MDA-MB-231 CRISPRi cells measured by CellTiter-Glo at 72 h treatment. IC50 (110 nM) is indicated above the curve. **C)** CRISPRi screen quality-control metrics showing sgRNA read-count evenness, missed sgRNAs, and mapping ratio at day 0 and day 18 of DMSO and THZ531 (110 nM) treatments. **D)** Distribution of gene-level β-scores for all and essential genes in DMSO- and THZ531 (110 nM)-treated CRISPRi screen arms. **E)** Read counts of individual sgRNAs targeting *ARID1A*, *SMARCE1*, and *SMARCB1*, and six randomly selected non-targeting sgRNAs, at the end of the screen in DMSO- and THZ531 (110 nM)-treated CRISPRi cells. **F)** Relative abundance of mCherry-positive CRISPRi cells targeting the indicated NuRD and NuRD-associated subunits during competition with GFP-positive sgRosa26 control cells under DMSO or THZ531 (110 nM) treatment for two weeks. sgRosa26-targeting cells are shown as a control. Data are shown as mean ± s.e.m. (n = 3).

**Figure S2.**
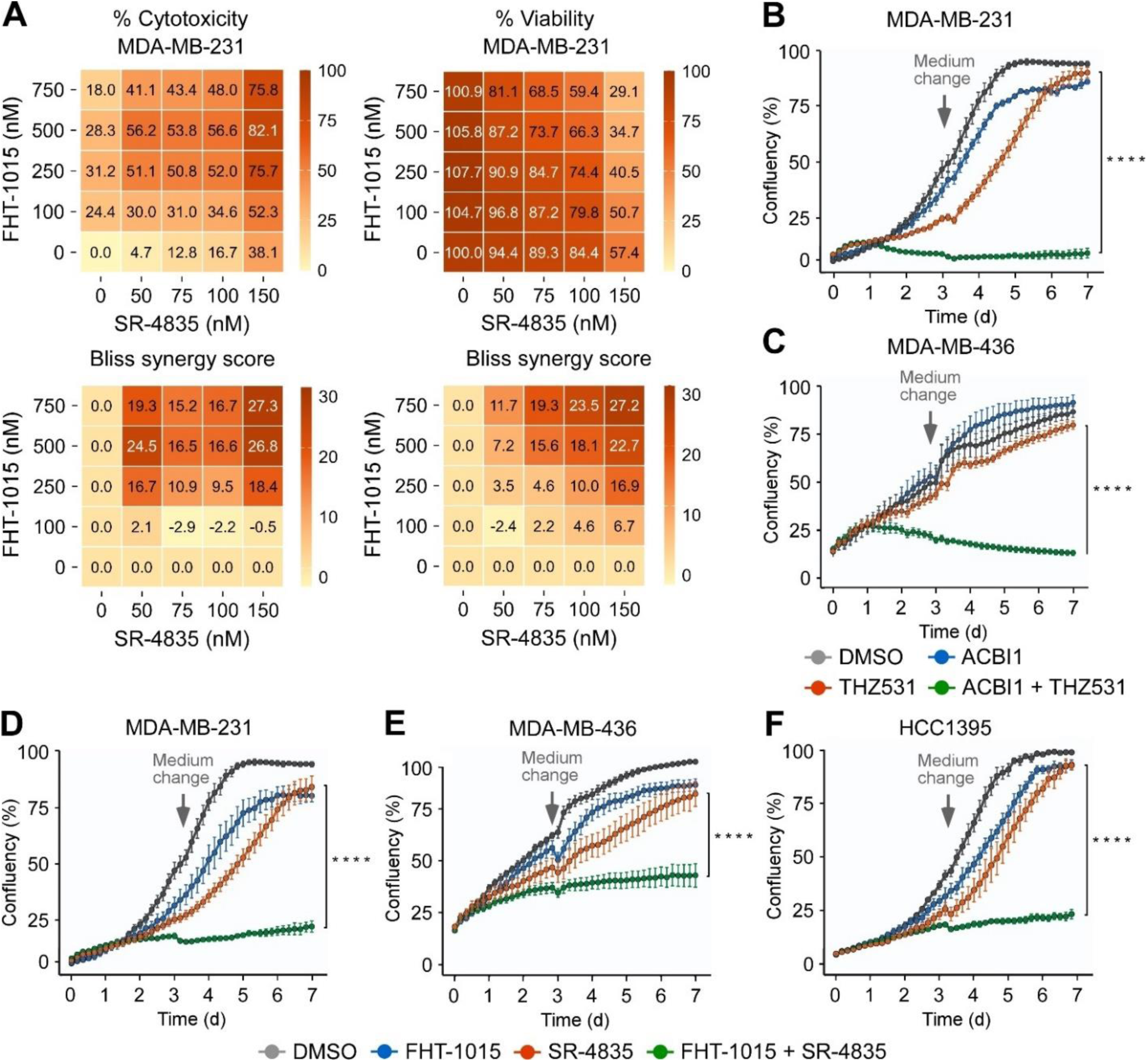
Pharmacological co-targeting of CDK12 and BAF synergizes in decreasing viability of TNBC models. **A)** Cytotoxicity and viability matrices with combinatorial titrations of FHT-1015 (A) and SR-4835 at indicated concentrations in MDA-MB-231 cells, depicting cytotoxicity and viability (top), and Bliss synergy scores (bottom) at 72 h. Cytotoxicity and viability data are shown as percentages relative to DMSO-treated cells and represent the average of independent experiments (n = 3). **B,C)** Cell growth of MDA-MB-231 (B) and MDA-MB-436 (C) cells treated for seven days with DMSO, ACBI1 (500 nM), THZ531 (60 nM), and the combination as indicated. Results are presented as % confluency and plotted as the mean ± s.e.m. (n = 3). Arrows indicate the replenishment of medium and drugs at day 3. \*\*\*\**P* < 0.0001, determined by two-way ANOVA using THZ531 and ACBI1 + THZ531 data sets. **D-F)** Cell growth of MDA-MB-231 (D), MD-MB-436 (E), and HCC1395 (F) cells treated for 7 days with FHT-1015 (D,E, 500 nM; F, 1 uM), SR-4835 (100 nM), and the combination as indicated. Results are presented as % confluency and plotted as the mean ± s.e.m. (n = 3). Arrows indicate the replenishment of medium and drugs at day 3. \*\*\*\**P* < 0.0001, determined by two-way ANOVA using SR-4835 and FHT-1015 + SR-4835 data sets.

**Figure S3.**
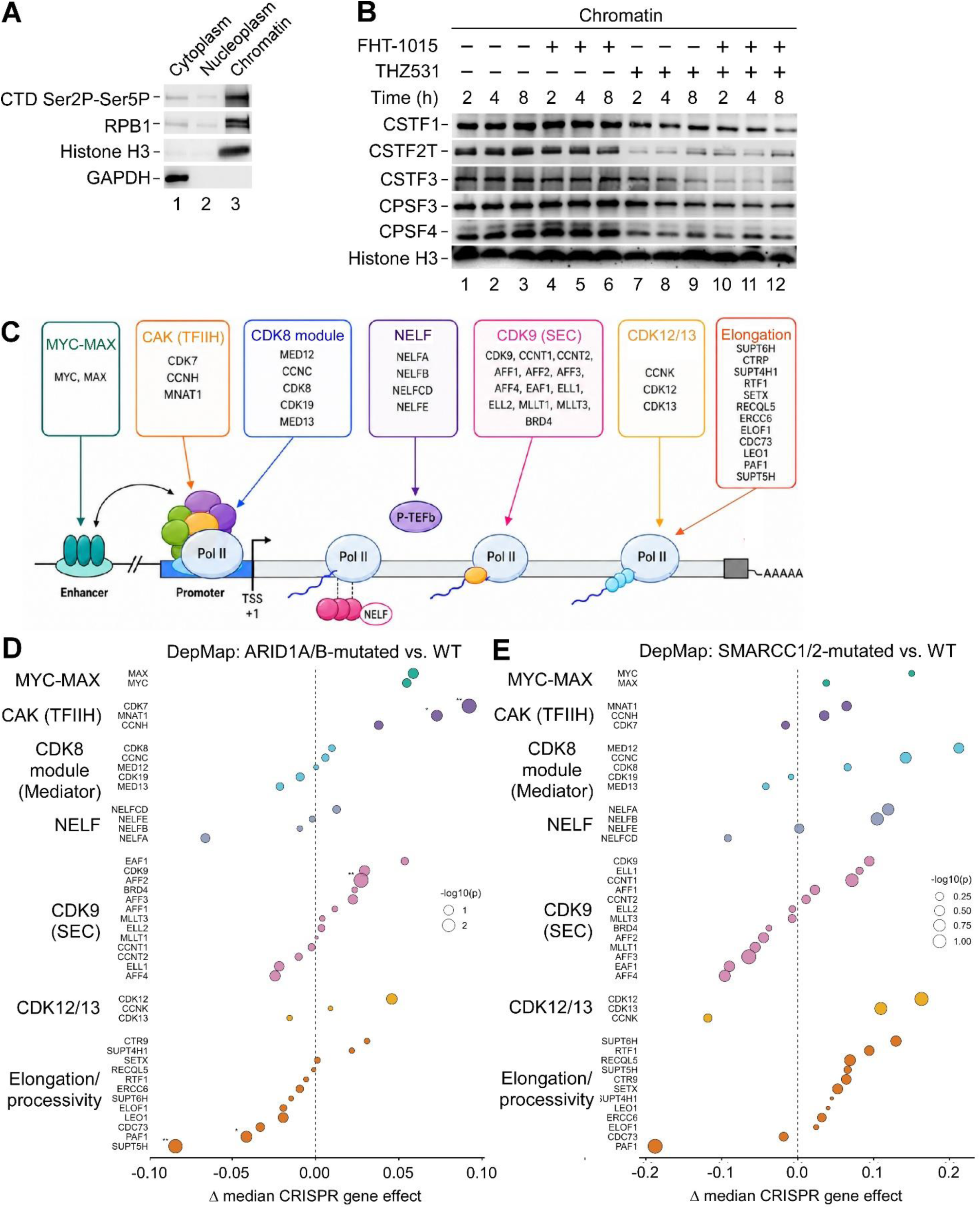
DepMap analysis of transcriptional dependency patterns in BAF-mutated cancer models. **A)** Immunoblot analysis of the indicated proteins in cytoplasmic, nucleoplasmic, and chromatin fractions of MDA-MB-231 cells. **B)** Immunoblot analysis of the indicated proteins in chromatin fractions of MDA-MB-231 cells treated with DMSO (–), FHT-1015 (750 nM), THZ531 (150 nM), and the combinations for 2, 4, and 8 h as indicated. **C)** Depiction of regulators controlling distinct steps of the Pol II transcription cycle that were used in DepMap CRISPR dependency analysis. **D)** DepMap CRISPR dependency analysis comparing ARID1A/B-mutated cancer models (n = 148) with the corresponding non-mutated models (n = 1044). Δ median gene-effect scores indicate differential dependency, with negative values reflecting stronger dependency in the BAF-mutated group. Point size indicates −log10(P); asterisks indicate nominal Wilcoxon rank-sum significance (*P < 0.05, **P < 0.01). **E)** DepMap CRISPR dependency analysis comparing SMARCC1/2-mutated cancer models (n = 15) with the corresponding non-mutated models (n = 1177). Δ median gene-effect scores indicate differential dependency, with negative values reflecting stronger dependency in the BAF-mutated group. Point size indicates −log10(P); asterisks indicate nominal Wilcoxon rank-sum significance (*P < 0.05, **P < 0.01).

**Figure S4.**
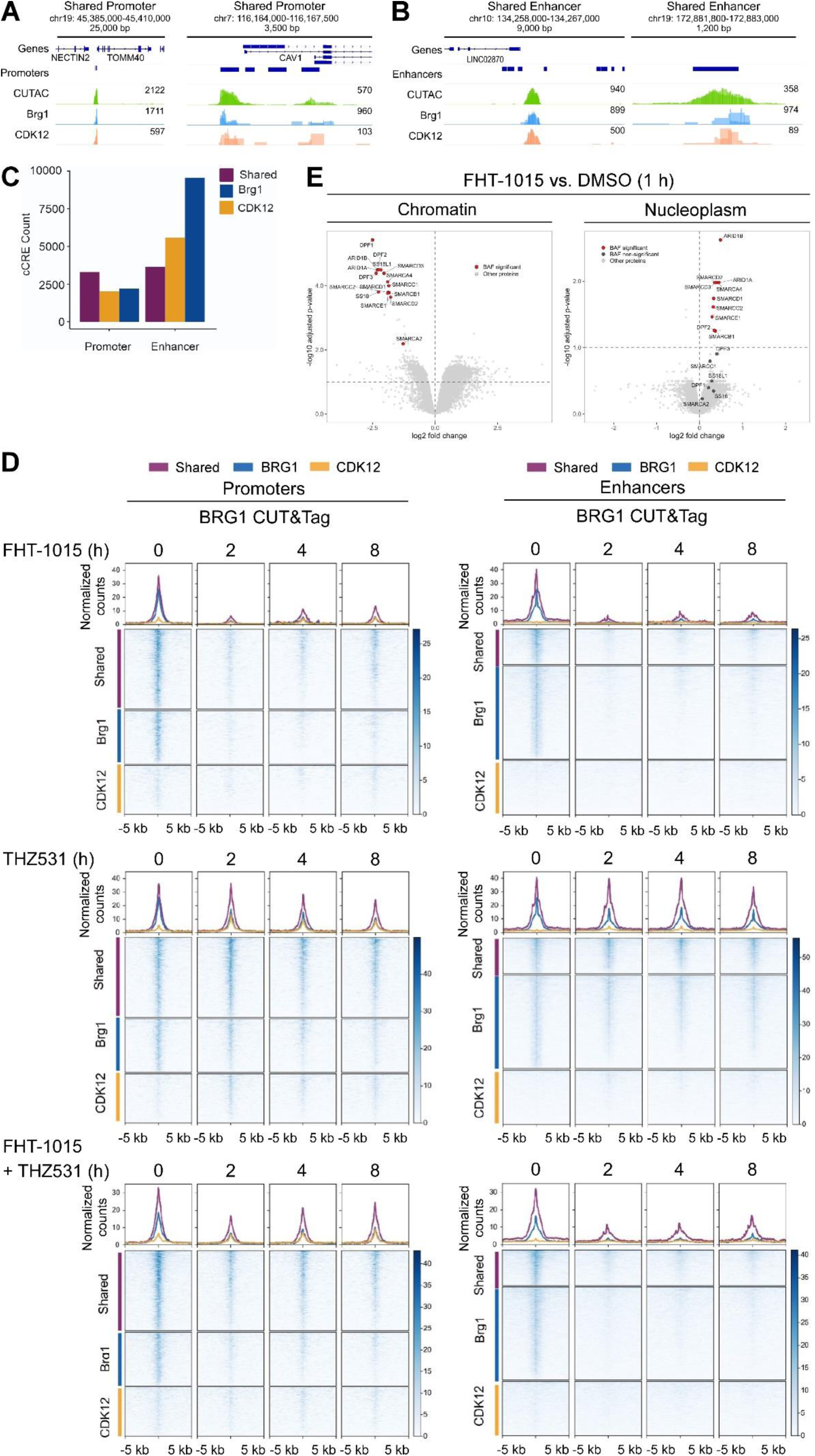
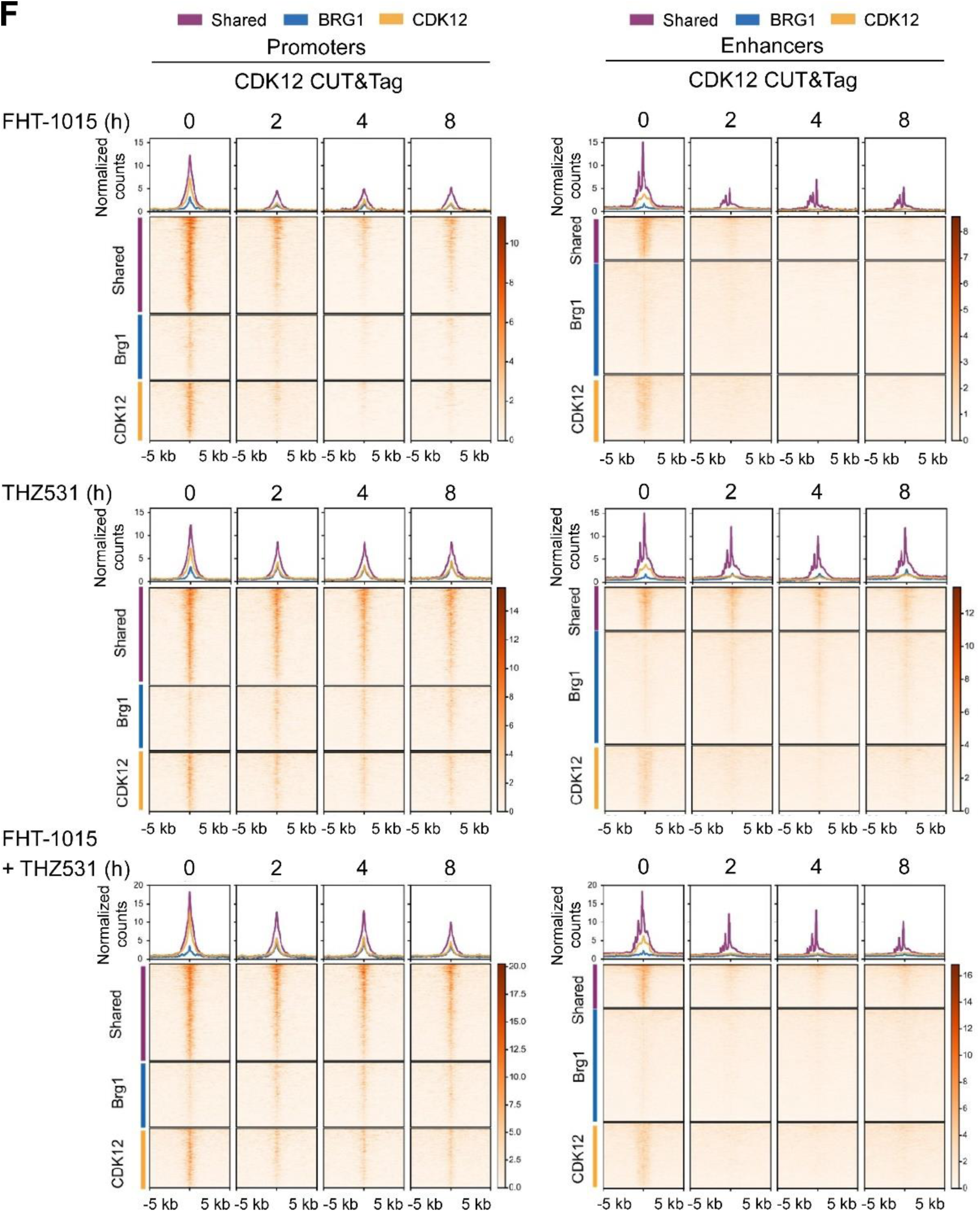
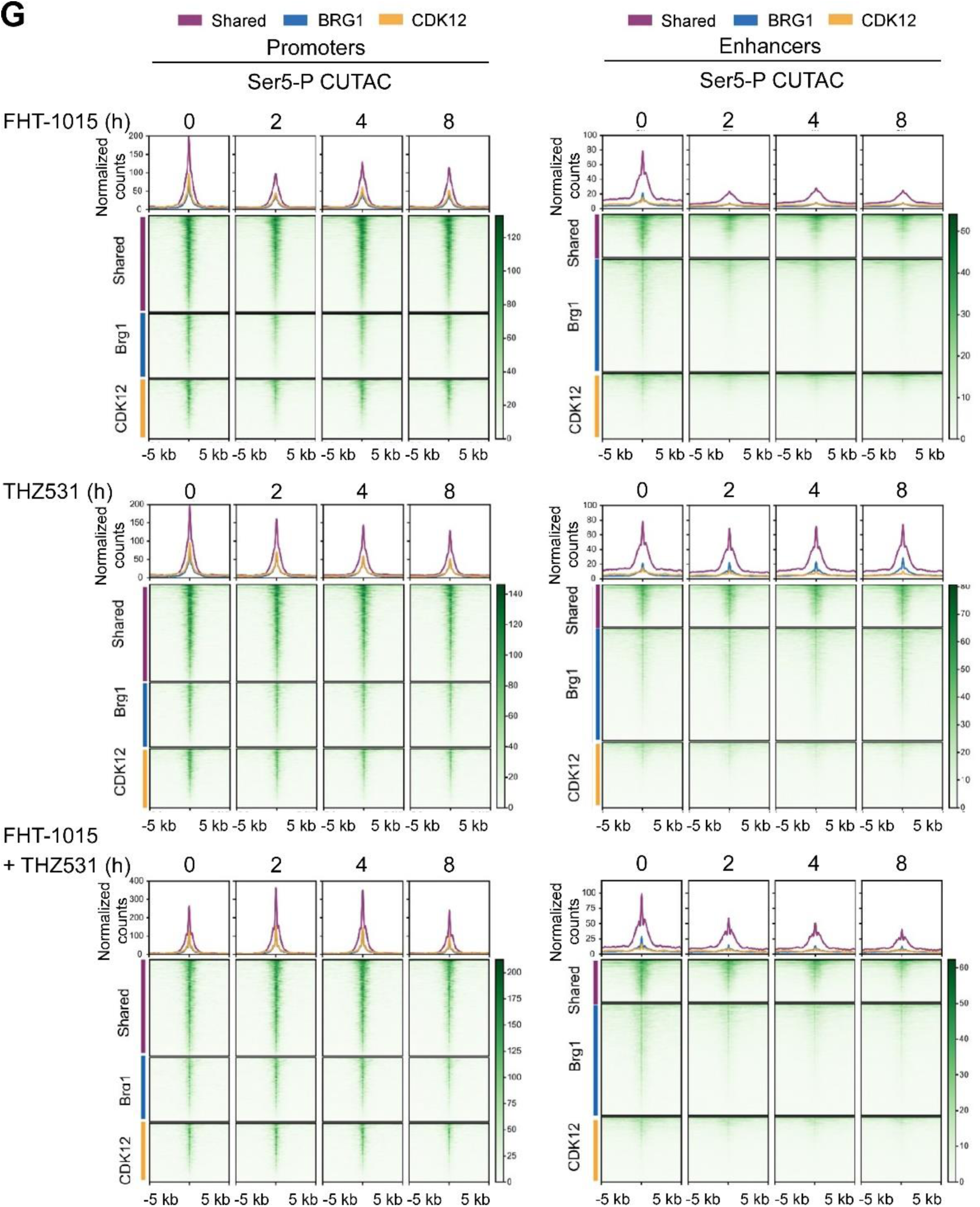
Catalytic activities of CDK12 and BAF promote their chromatin occupancy and transcription-coupled DNA accessibility. **A,B)** Representative genome browser tracks showing normalized Ser5-P CUTAC, BRG1 CUT&Tag, and CDK12 CUT&Tag signal at BRG1 and CDK12 co-occupied promoters (A) and enhancers (B). Track maxima are indicated on the right of each signal track. **C)** Number of BRG1 and CDK12 co-occupied (shared), BRG1-only, and CDK12-only promoters and enhancers identified by CUT&Tag. **D,F,G)** Average profiles (above) and heatmaps (below) of BRG1 CUT&Tag (D), CDK12 CUT&Tag (F), and Ser5-P CUTAC (G) signal at BRG1 and CDK12 co-occupied (shared), BRG1-only, and CDK12-only promoters and enhancers in MDA-MB-231 cells treated with FHT-1015 (1 μM), THZ531 (400 nM), and the combination for 2, 4, and 8 h as indicated (n = 2). Average profiles show normalized signal across the indicated regulatory-element classes. Promoter and enhancer profiles are centered on transcription start sites and enhancer mid-points, respectively. **E)** Volcano plots showing log2FC values (adjusted *P* ≤ 0.1) of BAF subunit levels in chromatin and nucleoplasmic fractions of MDA-MB-231 cells treated with FHT-1015 (1 μM) for 1 h, relative to DMSO (n = 4). Significant and non-significant BAF subunits are shown in red and dark gray, respectively; other proteins are shown in light gray.

**Figure S5.**
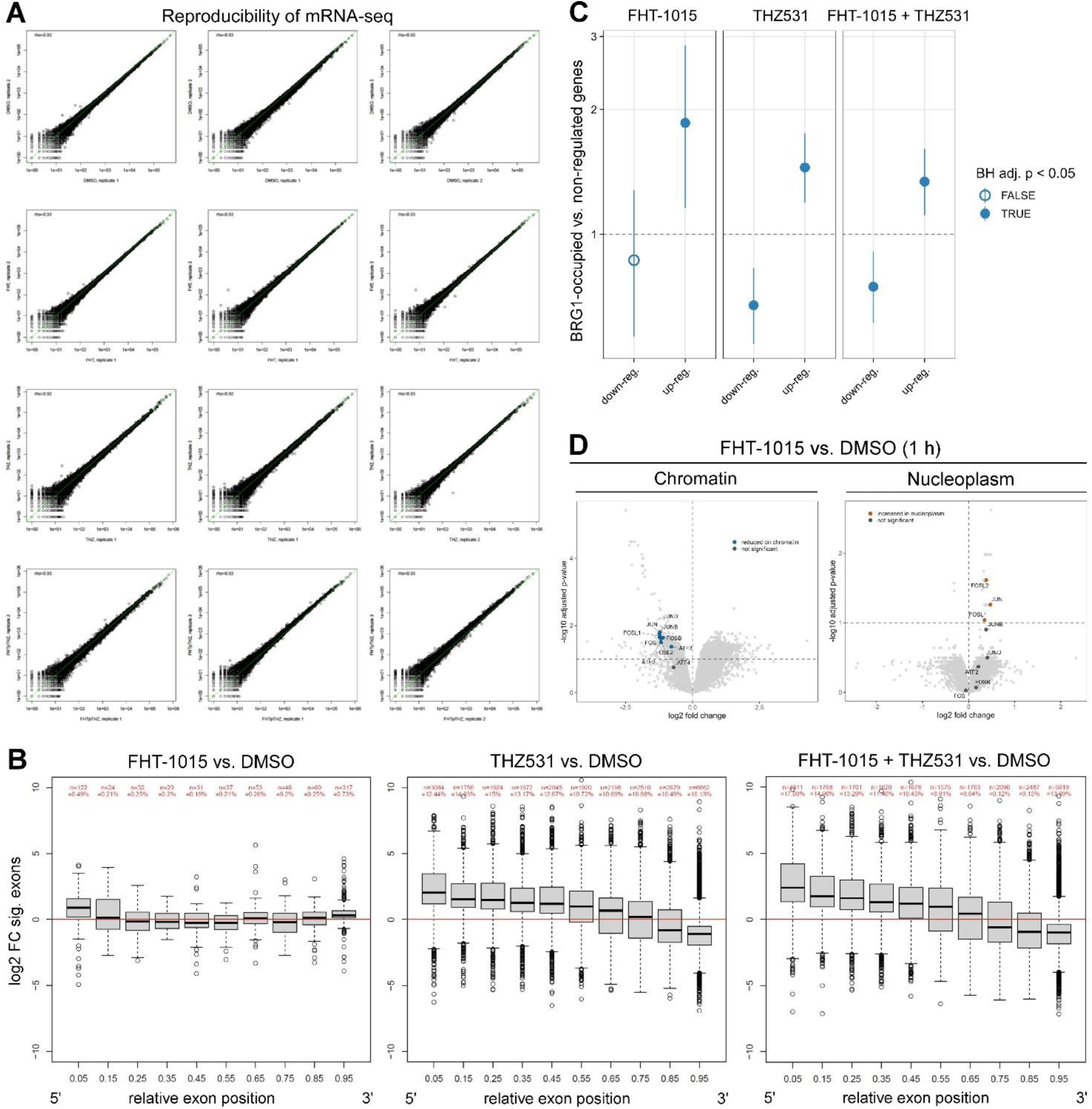
CDK12 and BAF perturbation reshapes gene expression programs in TNBC cells. **A)** Pairwise scatterplots comparing raw read counts between mRNA-seq replicates (n = 3). Spearman correlation coefficients (ρ) are indicated. **B)** Differential exon-usage analysis showing exon-level log2FC changes for differentially used exons (adjusted P ≤ 0.001) across relative exon positions within genes of MDA-MB-231 cells treated for 8 h with DMSO (–), FHT-1015 (750 nM), THZ531 (300 nM), and the combination as indicated, relative to DMSO (n = 3). Exons were binned into 10 equal-sized bins for each gene. Numbers on top indicate the number and percentage of differentially used exons in each bin. **C)** Odds-ratio analysis testing enrichment of BRG1 promoter occupancy among RNA-seq-regulated genes relative to non-regulated genes after the indicated 8 h treatments. BRG1-occupied promoters include BRG1-only and BRG1/CDK12 co-occupied promoters. Points show odds ratios from Fisher’s exact tests with 95% confidence intervals; filled circles indicate BH-adjusted P < 0.05. **D)** Volcano plots showing log2FC values (adjusted *P* ≤ 0.1) of AP-1 subunit levels in chromatin and nucleoplasmic (bottom) fractions of MDA-MB-231 cells treated with FHT-1015 (1 μM) for 1 h, relative to DMSO (n = 4). Significant and non-significant AP-1 subunits are shown in red and dark gray, respectively; other proteins are shown in light gray.

**Figure S6.**
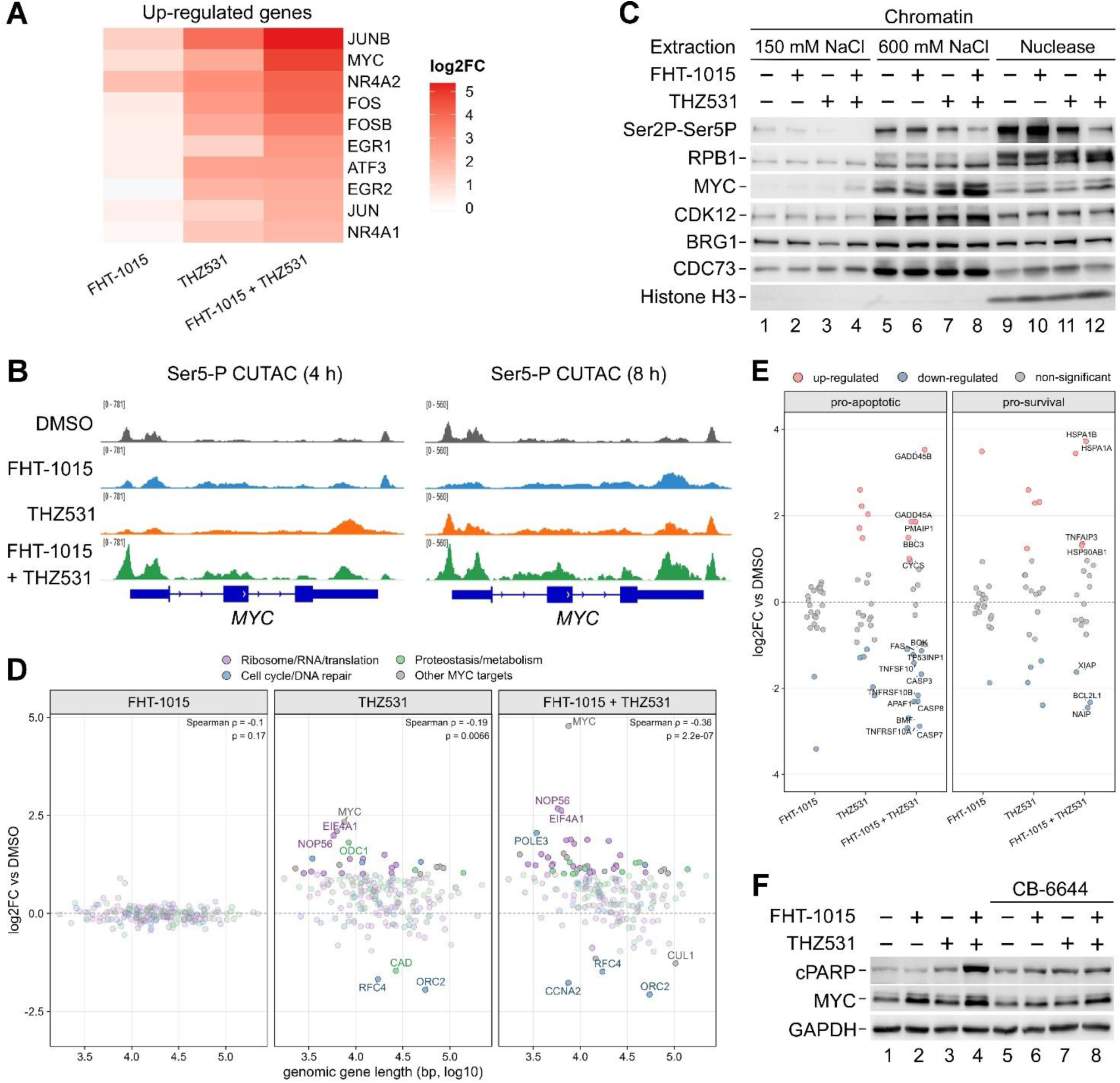
MYC-linked transcriptional and apoptotic responses to CDK12–BAF co-inhibition. **A)** Heatmap showing mRNA-seq log2FC values (adjusted P ≤ 0.001) of short, immediate-early transcriptional regulators of MDA-MB-231 cells treated for 8 h with DMSO (–), FHT-1015 (750 nM), THZ531 (300 nM), and the combination as indicated, relative to DMSO (n = 3). **B)** Ser5-P CUTAC genome browser tracks at the *MYC* locus of MDA-MB-231 cells treated for 4 h and 8 h with DMSO, FHT-1015 (1 μM), THZ531 (400 nM), and the combination as indicated. Each track shows a representative Ser5-P CUTAC replicate (n = 2). **C)** Immunoblot analysis of the indicated proteins upon sequential salt and Nuclease extraction of chromatin fractions of MDA-MB-231 cells treated for 24 h with DMSO (–), FHT-1015 (750 nM), THZ531 (150 nM), and the combination as indicated. **D)** mRNA-seq log2FC values of HALLMARK_MYC_TARGETS_V1 genes plotted by genomic gene length of MDA-MB-231 cells treated for 8 h with DMSO, FHT-1015 (750 nM), THZ531 (300 nM), and the combination as indicated (n = 3). Significantly regulated genes depicted as circles are colored by curated functional categories, and select genes are labeled. Spearman correlation coefficients and P values are indicated. **E)** mRNA-seq log2FC values DMSO (adjusted P ≤ 0.001) of curated pro-apoptotic and pro-survival genes of MDA-MB-231 cells treated for 8 h with FHT-1015 (750 nM), THZ531 (300 nM), and the combination as indicated, relative to DMSO (n = 3). Red and blue circles depict significantly up-regulated and down-regulated genes, respectively; gray circles indicate non-significantly regulated genes. Genes regulated upon the combination treatment are labeled. **F)** Immunoblot analysis of the indicated proteins in whole cell extracts of MDA-MB-231 cells treated for 24 h with DMSO (–), FHT-1015 (750 nM), THZ531 (150 nM), and the combination with or without CB-6644 (200 nM) as indicated.

**Figure S7.**
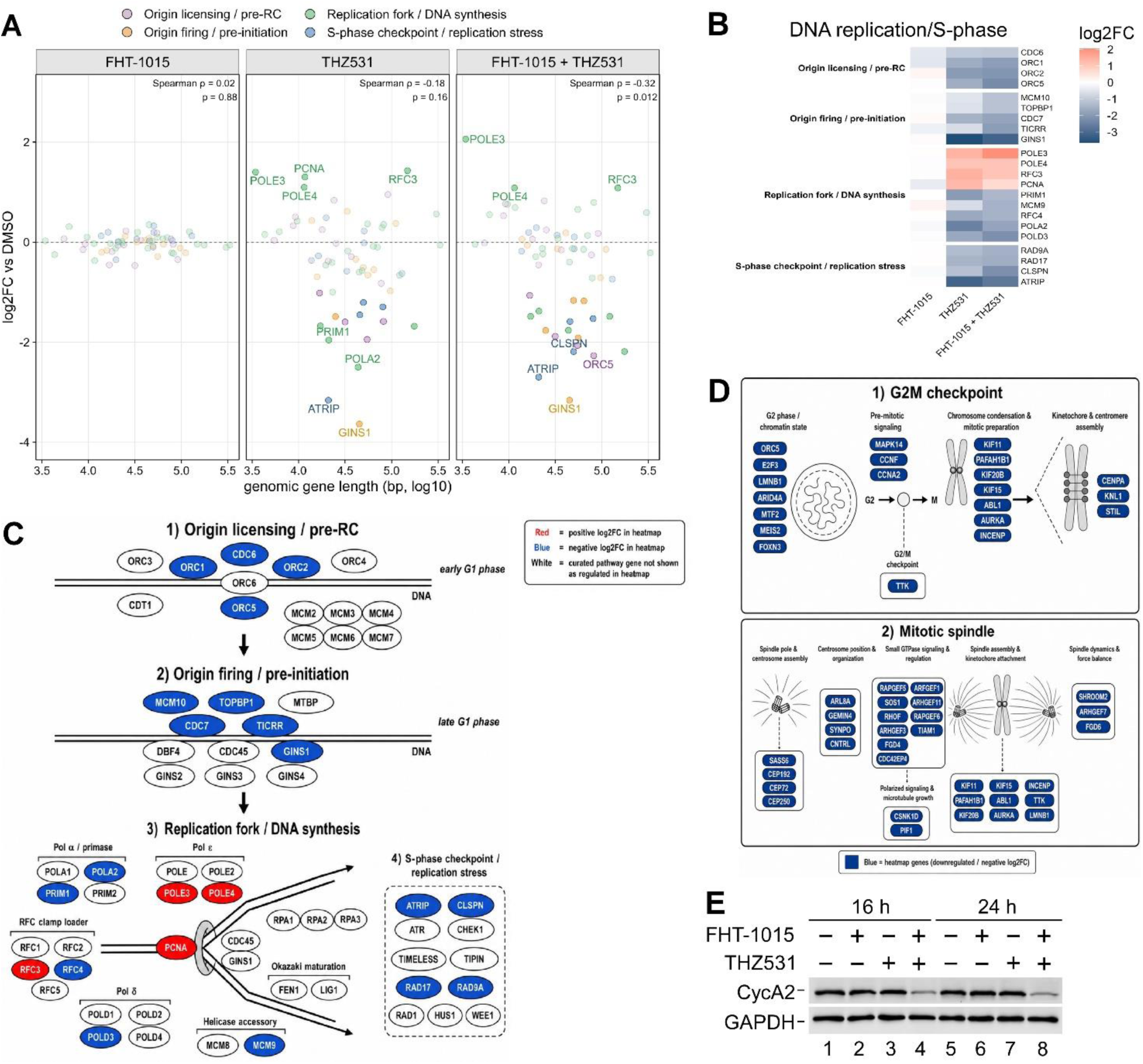
RNA-seq analysis of DNA replication and mitotic checkpoint gene programs. **A)** mRNA-seq log2FC values of curated DNA replication and S-phase progression genes plotted by genomic gene length of MDA-MB-231 cells treated for 8 h with FHT-1015 (750 nM), THZ531 (300 nM), and the combination as indicated, relative to DMSO (n = 3). Significantly regulated genes depicted as circles are colored by replication function categories, and select genes with the largest absolute log2FC values are labeled. Spearman correlation coefficients and P values are indicated. **B)** Heatmap showing mRNA-seq log2FC values (adjusted P ≤ 0.001) of curated DNA replication/S-phase genes of MDA-MB-231 cells treated for 8 h with FHT-1015 (750 nM), THZ531 (300 nM), and the combination as indicated, relative to DMSO (n = 3). **D)** Schematic of curated DNA replication/S-phase genes shown in (B) mapped to replication licensing, origin firing, fork progression, and S-phase checkpoint modules. Node colour indicates RNA-seq log2FC: blue, down-regulated; red, up-regulated; white, non-significantly regulated genes. **E)** Schematic of non-additively down-regulated G2/M checkpoint and mitotic spindle genes shown in Fig. 7D. Blue nodes indicate genes more strongly repressed by the combined FHT-1015 + THZ531 treatment than by either inhibitor alone. **F)** Immunoblot analysis of the indicated proteins in whole cell extracts of MDA-MB-231 cells treated for 16 h and 24 h with DMSO (–), FHT-1015 (750 nM), THZ531 (150 nM), and the combination as indicated.

## Notes

### Competing Interest Statement

The authors have declared no competing interest.

### Summary of Updates

The revised manuscript includes new results presented in Figures 2 and 3.

